# O-GalNAc glycans enrich in white matter tracts and regulate nodes of Ranvier

**DOI:** 10.1101/2024.05.09.593410

**Authors:** Maxence Noel, Suttipong Suttapitugsakul, Richard D. Cummings, Robert G. Mealer

**Affiliations:** Department of Surgery, Beth Israel Deaconess Medical Center, Harvard Medical School, Boston, MA 02215, USA; Department of Psychiatry, Oregon Health & Science University, Portland, OR 97239

**Keywords:** Mammalian brain, Glycosylation, O-glycosylation, O-GalNAc, Cosmc, white matter, node of Ranvier, Siglec-4, lecticans, cell adhesion

## Abstract

Protein O-glycosylation is a critical modification in the brain, as genetic variants in the pathway are associated with both common and severe neuropsychiatric phenotypes. However, little is known about the most abundant type of O-glycan in the mammalian brain, which are O-GalNAc linked. Here, we determined the spatial localization, protein carriers, and cellular function of O-GalNAc glycans in mouse brain. We observed striking spatial enrichment of O-GalNAc glycans in white matter tracts and at nodes of Ranvier. Glycoproteomic analysis revealed that more than half of the identified O-GalNAc glycans were present on chondroitin sulfate proteoglycans termed lecticans, and display both domain enrichment and site-specific heterogeneity. Though genetic ablation in a single cell type failed to replicate the severe phenotypes seen in congenital disorders, inhibition of O-GalNAc synthesis in neurons reduced binding of Siglec-4, a known regulator neurite growth, and shortened the length of nodes of Ranvier. This work highlights a new function of O-GalNAc glycans in the brain and will inform future studies on their role in development and disease.

## Introduction

Glycosylation is the enzymatic attachment of carbohydrates to proteins and lipids, which generates an enormous diversity of non-template-based biomolecules involved in health and disease^1^. In the brain, glycosylation is key for proper development and function, affecting nearly all processes including synaptic plasticity^2^, ion channel function^3^, and axonal guidance^4^. Severe mutations in glycosylation genes result in congenital disorders of glycosylation (CDGs) which frequently involve the brain, as over 80% of CDG cases include neurological dysfunction^5^. The role of abnormal glycosylation in common disorders of the brain is becoming increasingly apparent both from genetic associations^6–8^ and post-mortem studies^9,10^.

In the brain, most studies on glycosylation have focused on glycolipids^11^ and asparagine linked (N-) glycoproteins^12,13^. Less is known about glycans linked to serine, threonine, and occasionally tyrosine residues, termed O-glycoproteins, which include at least 10 independent pathways utilizing distinct enzymes and carbohydrate substrates^14^. Two exceptions are O-mannose (O-Man) glycans and O-N-acetylglucosamine (O-GlcNAc) glycans. Mutations in genes encoding the synthetic enzymes of the POMT-initiated O-Man pathway can lead to congenital muscular dystrophies^15^; O-GlcNAc glycans, which are present on nuclear and cytoplasmic proteins, are implicated in aging and neurodegeneration^16^. However, in recent studies using different methodologies for glycan release and analysis, 70-80% of all brain O-glycans were of the N-acetylgalactosamine (O-GalNAc), or mucin-type (here termed O-glycans)^17,18^.

In contrast to most O-glycosylation pathways beginning in the endoplasmic reticulum (ER) or the O-GlcNAc nuclear/cytosolic pathway, O-GalNAc glycosylation is initiated in the Golgi apparatus by a family of 20 isoenzymes termed N-acetylgalactosaminyl-transferases, or GALNTs, generating a unique monosaccharide epitope termed Tn antigen (GalNAcα1-Ser/Thr)^14,19^. In normal tissues, Tn antigen is efficiently modified with galactose by the enzyme C1GALT1, or T-synthase, in complex with its exclusive and essential chaperone C1GALT1C1, or Cosmc, to generate the disaccharide structure termed T antigen, or Core 1 (Galβ1-3GalNAcα1-Ser/Thr). The T antigen can then be modified by different glycosyltransferases to generate an array of complex and extended O-glycans (**Fig 1A**), which are commonly illustrated using the SNFG nomenclature (**Box 1**)^20,21^. Across the mammalian brain, we previously demonstrated that nearly 80% of O-glycan structures are extensions of the T antigen and contain sialic acid^18^.

**Figure 1.**
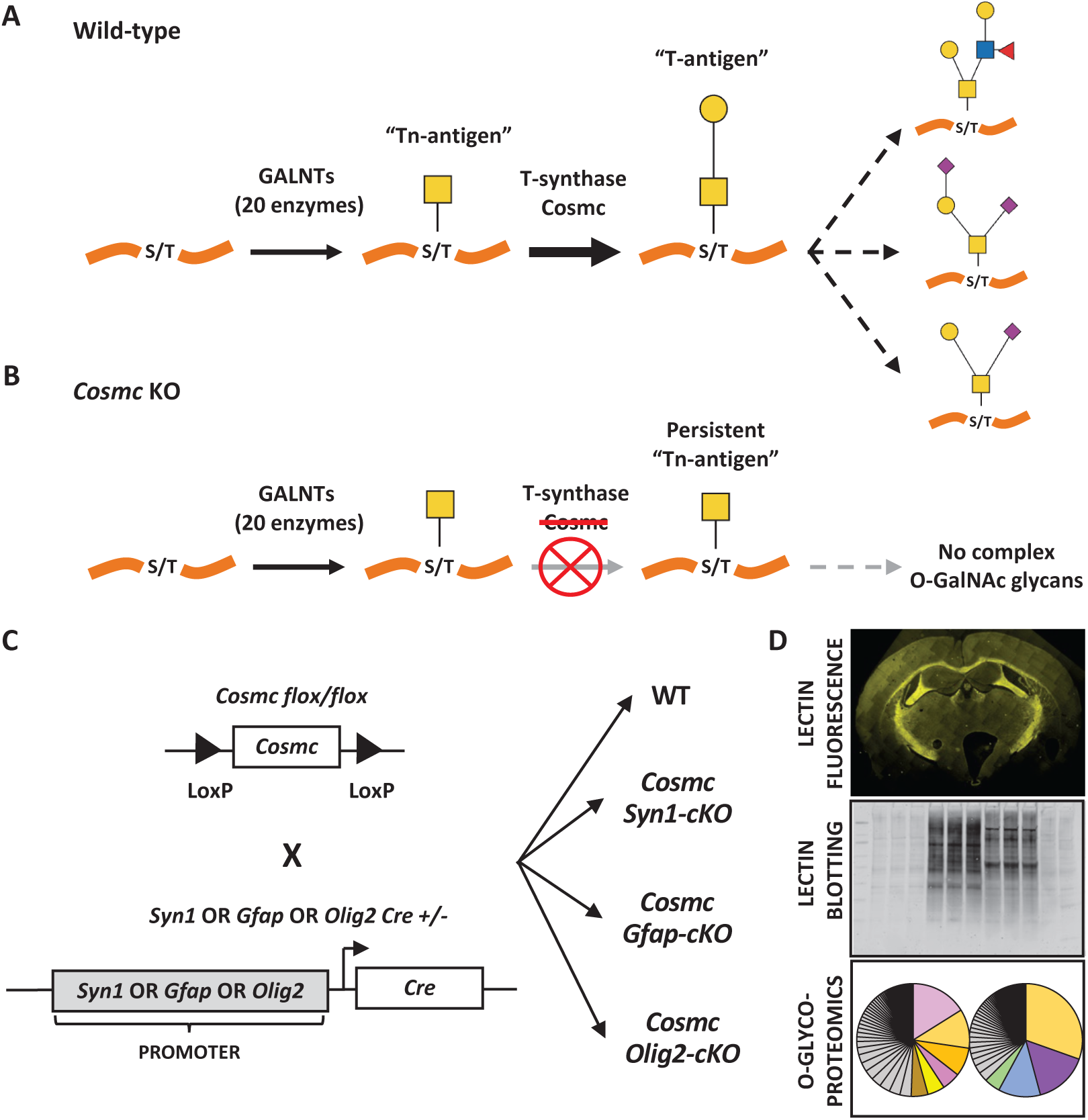
Targeting mucin-type O-GalNAc glycosylation in the brain. **A)** Under basal conditions, O-GalNAc glycosylation is commonly initiated on serine and threonine residues by a family of 20 isoenzymes termed GALNTs to generate “Tn-antigen.” This structure is extended by T-synthase, which requires its exclusive chaperone Cosmc, to generate the “T-antigen,” which can then be modified by multiple enzymes to generate the diverse pool of complex O-GalNAc glycans observed in tissues. **B)** Genetic deletion of Cosmc inhibits T-synthase activity, resulting in persistent expression of “Tn-antigen” and a lack of all complex O-GalNAc glycans. **C)** Experimental plan to study the role of O-GalNAc glycans in brain: female floxed *Cosme* mice were bred with male *Synl*-Cre, *Gfap*-Cre, or *O/ig2*-Cre mice, to generate brain tissue for analysis. **D)** Parallel analyses using lectin fluorescence, lectin blotting, and O-glycoproteomics were used to understand to role of O-GalNAc glycosylation in the brain.

**Box 1.**
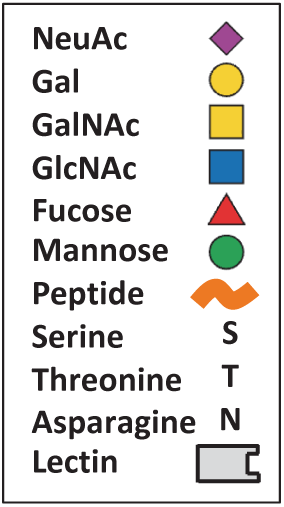
Key according to Symbol Nomenclature for Glycans (SNFG), amino acid code, and lectin schematics.

Complex O-glycans are essential for life, as global genetic deletion of either T-synthase or Cosmc prevents O-GalNAc elongation (**Fig. 1B**) and causes early embryonic lethality in mice^22,23^. Tissue and cell-specific roles of O-glycans have been revealed using targeted deletion of Cosmc or T-synthase, and implicate O-glycans in biological processes including regulation of gut microbiota^24^, platelet biogenesis^25^, lymphangiogenesis^26^, podocyte functions^27^ and immunity^28–30^. Few studies address O-glycans in mammalian brain, despite heritable conditions supporting their importance for proper function and development. Genes for at least two GALNTs (*GALNT10* and *GALNT17*, also known as *WBSCR17*) are significantly associated with schizophrenia by GWAS^31^. *GALNT2* mutations in humans cause a CDG presenting with severe neuropsychiatric symptoms including intellectual disability, seizures, and structural abnormalities, and was mirrored in rodent models of *Galnt2* mutation^32^. Changes in Cosmc/T-synthase levels have been reported in Alzheimer’s disease^33^ and our group in collaboration with others recently described an X-linked CDG including intellectual disability caused by *COSMC* (*C1GALT1C1*) mutation in human male carriers^34^.

However, little is known about the functions and glycoprotein expression of O-GalNAc glycans in the mammalian brain. Studies in drosophila have shown that O-glycans are essential in brain development, morphology, and synaptic function^35^. However, the bulk of drosophila brain O-glycans are unsialylated Tn antigen and T antigen, which are nearly undetectable in the mammalian brain at baseline^18^. Here, we performed targeted genetic blockade of the O-GalNAc pathway (**Fig. 1C**) to characterize the spatial localization, glycoproteomic landscape, and functional relevance of O-GalNAc glycans in mammalian brain (**Fig. 1D**). Our results support an unexpected role for O-glycans in white matter tracts and at nodes of Ranvier, with their synthesis and function being implicated in development and disease.

## Results

### Biosynthetic enzymes for complex O-glycans are expressed across all brain cell types

To predict which cell types are capable of synthesizing O-glycans, we downloaded single cell expression data from mouse cortex^36^ and compared levels of transcripts for the relevant glycosyltransferases and chaperones for the first two steps in the pathway (**Supplemental Table 1**). Although expression levels in the brain are low, consistent broadly with all glycosylation genes in the brain^18^, transcripts for many of the initiating *Galnt* family members, the extending *T-synthase* (*C1galt1*), and its chaperone *Cosmc* (*C1galt1c1*), were present across all cell classes, suggesting each possess the enzymes necessary to synthesize complex O-glycans.

### Endogenous O-glycans are enriched in white matter tracts

We sought to directly visualize the localization of endogenous O-glycans in the wildtype brain using the lectin peanut agglutinin (*Arachis hypogaea*) (PNA), which recognizes the terminal galactose of the T-antigen disaccharide in O-glycans^37^. However, PNA will not bind well to those that are sialylated (**Fig. 2A**)^38^; we previously observed that >90% of O-GalNAc glycans in the murine brain are sialylated^18^. Baseline PNA distribution in the wild-type cerebellum showed a striking pattern of enrichment within white matter tracts of the arbor vitae (**Fig. 2B**). Treatment with neuraminidase (NeuA) to remove sialic acid resulted in a robust elevation of PNA binding across the cerebellum with the most intense signal remaining in white matter tracts. Co-treatment with O-glycosidase and NeuA, which specifically releases the T antigen (Gal-GalNAc-S/T)^39^, caused a dramatic reduction of PNA binding in most areas including white matter tracts, though some signal persisted that appeared to be O-glycosidase insensitive within a pattern resembling microvasculature (**Supplemental Fig. 1**). This may result from glycans that were physically inaccessible to the enzyme or have another modification that does not affect PNA binding but blocks O-glycosidase effectiveness. The presence of 200 mM galactose blocked PNA binding, consistent with specific PNA binding to terminal galactose of O-glycans (**Fig. 2B**). Binding of the lectin Jacalin, which is reported to have an affinity for terminal galactose of O-glycans independent of sialic acid^40^, showed a diffuse pattern across each layer of the cerebellum (**Supplemental Fig. 2**). However, treatment with O-glycosidase and NeuA only reduced Jacalin binding in the arbor vitae, consistent with the enrichment of O-glycans to white matter tracks. The presence of 800 mM galactose blocked nearly all Jacalin binding including in the arbor vitae (**Supplemental Fig. 2**).

**Figure 2.**
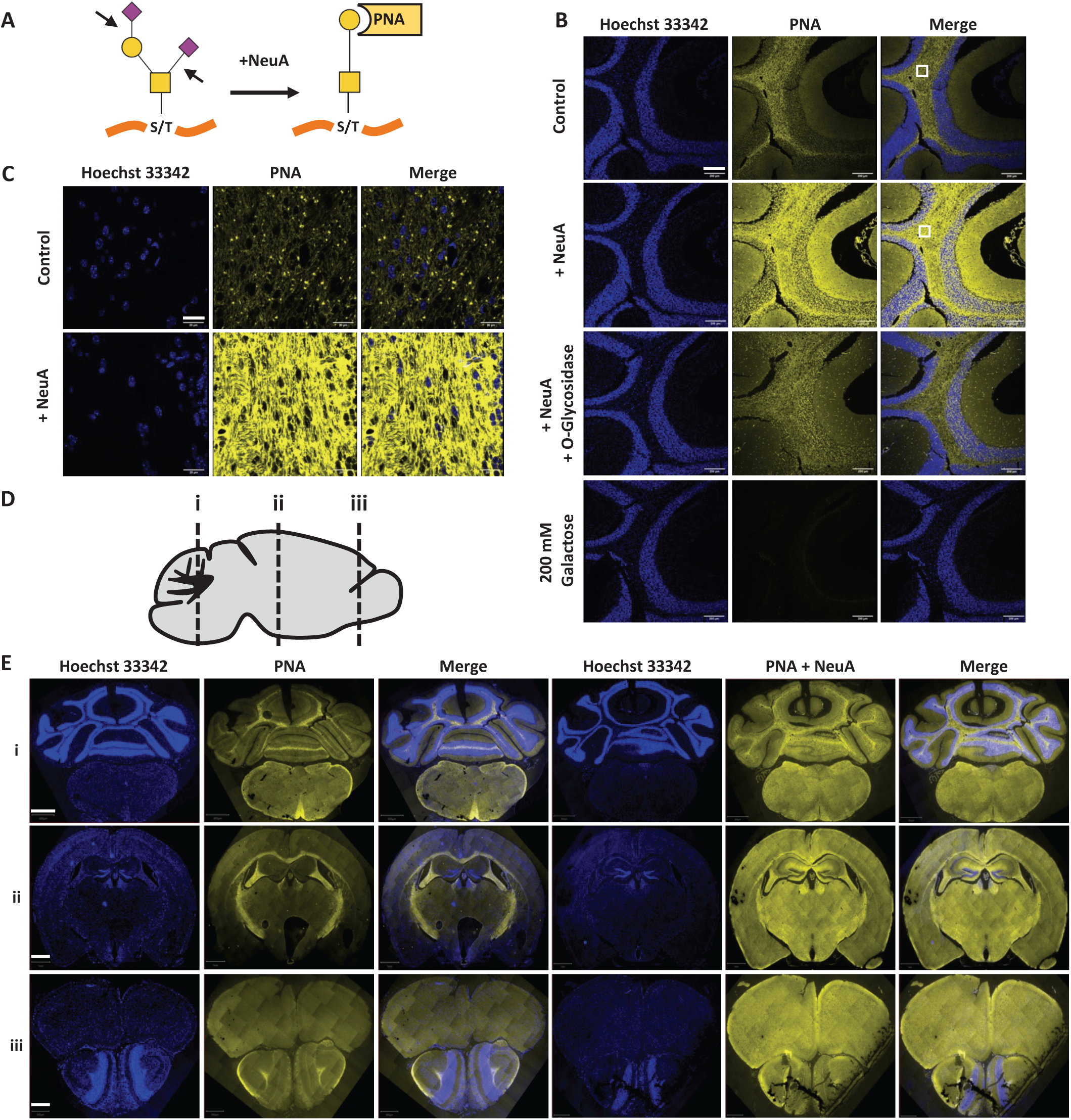
PNA binding enriches in white matter tracts. **A)** PNA preferentially binds galactose of O-GalNAc glycans lacking sialic acid. **B)** PNA binding increases in brain sections following neuraminidase treatment, is sensitive to O-glycosidase, and is blocked by galactose. Scale bar = 200 µm. White square represents ROI for (C). **C)** High magnification of PNA binding in white matter from ROI (white square) in (B). Scale bar = 20 µm. **D)** Schematic illustrating the location of coronal sections for (E). **E)** PNA binding across multiple brain regions before and after neuraminidase treatment. Scale bars i = 800 µm, ii = 1 mm, iii = 500 µm. Images with additional detail and annotations are included in Supplementary material.

High magnification of PNA binding in the arbor vitae showed several dense puncta with more diffuse signal after NeuA pre-treatment (**Fig. 2C**, **Supplemental Fig. 3**). Coronal sections across multiple brain regions (**Fig. 2D**) showed consistent PNA staining in white matter tracts (**Fig. 2E**, **Supplemental Fig. 4**). In the most caudal section, PNA staining was most prominent in the arbor vitae of the cerebellum and pyramid/corticospinal tract of the brain stem. In the middle section, prominent PNA staining was observed in the corpus callosum, fimbria, and internal capsule. In the most rostral section, diffuse staining was seen throughout the frontal cortex with increased signal in the anterior commissure and lateral tract of the olfactory bulb. A similar pattern of increased binding in white matter tracts was observed with PNA following NeuA (**Fig. 2E**). The punctate pattern of PNA binding observed in the arbor vitae of the cerebellum (**Fig. 2C**, **Supplemental Fig. 3**) was also present in the olfactory bulb, where the trajectory of white matter tracts would be orthogonal to the coronal section (**Supplemental Fig. 5**). In contrast, the pattern of PNA binding observed in the corpus callosum, where white matter tracts run parallel to the coronal section, appeared striated both with and without NeuA pre-treatment (**Supplemental Fig. 5**).

### Brain O-glycans display a distinct spatial distribution compared to N-glycans

The unexpected enrichment of O-glycans in white matter tracts led us to compare their spatial distribution to that of N-glycans in the brain. We focused on the cerebellum due to its distinct layering and compartmentation, including the synapse rich molecular layer (ML), the granular layer (GL) with many cell bodies/nuclei, and the arbor vitae (AV) containing axons surrounded by myelin. As previously described^41^, lectins specific for N-glycans (ConA, GNL, AAL, PHA-E and RCA-I) all showed predominant binding for the ML, with differing levels of binding seen for the GL and AV (**Fig. 3A**). PNA signal in the brain was unique in its enrichment for white matter tracts including the AV both with or without NeuA pre-treatment (**Fig. 3A**). Quantification of the relative signal intensity across the cerebellum for multiple lectins shows that while each displays a distinct pattern, only PNA has the highest affinity for the AV, both with and without NeuA pre-treatment, and highlights the different spatial expression of N- and O-glycans (**Fig. 3B**).

**Figure 3.**
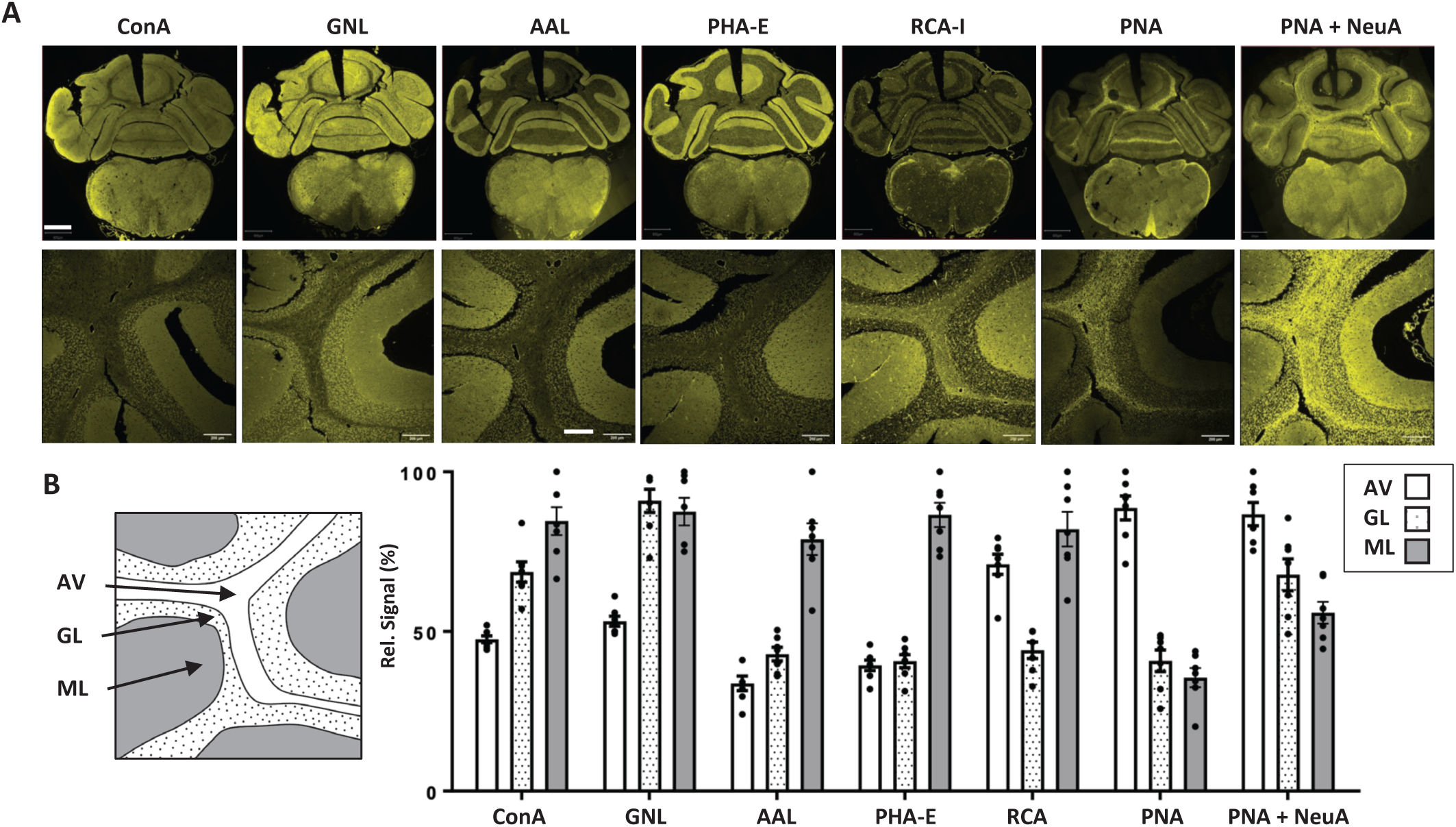
O-GalNAc enrichment in white matter tracts is distinct from N-glycans. **A)** Lectin binding in the cerebellum. Top row, cerebellum and brain stem. Scale bar = 800 µm. Bottom row, 20 X cerebellum. Scale bar = 200 µm. **B)** Schematic illustrating different layers of the cerebellum at 20X including the arbor vitae (AV), granular layer (GL), and molecular layer (ML) and relative quantification of lectin binding across the layers of the cerebellum from (A). Data presented as the mean binding intensity normalized to the highest value +/- SEM, with individual data points from at least six different cerebellar locations represented.

### Inhibition of O-glycan extension in neurons or astrocytes produced no overt phenotype

To prevent O-GalNAc extension in the brain by the T-synthase and avoid the early embryonic lethality associated with global deletion^22,23^, we first crossed floxed-*Cosmc* females (*Cosmc^f/f^*)^22^ with males from the well characterized driver lines *Syn1-Cre*^+42^, and *Gfap-Cre*^+43^ (**Fig. 1C**). *Syn1* promoter was used to target the deletion to neurons, though its expression in the mouse brain is broad including most neurons and many other cells, whereas *Gfap* expression is restricted primarily to astrocytes (**Supplemental Table 1**). Based on this pattern we predicted a large loss of extended O-glycans in *Cosmc^f/y^ Syn1-Cre^+^* conditional knockouts (*Cosmc Syn1-cKO*) and a narrow loss in *Cosmc^f/y^ Gfap-Cre^+^* conditional knockouts (*Cosmc Gfap-cKO*). As *Cosmc* is an X-linked gene with an associated X-linked disorder^34^, this breeding strategy produced wild-type and knock-out males as well as wild-type and mosaic females.

While the mosaic females are not a target of our study here, they have proven useful as an intermediate mosaic phenotype for *Cosmc* deletion when the full knock-out is too severe for analysis^22,24^. Birth ratios between male and females were similar in both crosses, and though there were slightly fewer *Cosmc Syn1-cKO* males and *Cosmc Syn1-cKO* mosaic females compared to WTs, it was not statistically significant (**Supplemental Table 2**). No early postnatal lethality or gross behavioral/neurological abnormalities were observed in either cKO line through early adulthood. These results suggested that inhibited expression of extended O-glycans in neurons and astrocytes did not result in early lethality or gross pathology, and that more subtle perturbations would be associated with changes in O-GalNAc synthesis.

### O-glycan levels were unchanged by *Cosme* deletion in neurons and astrocytes

In the absence of an overt neurological phenotype, we sought to confirm the biochemical inhibition of O-glycan extension in the brain prior to additional analyses. Blocking O-glycan elongation by Cosmc/T-synthase preserves GALNT-mediated initiation on native protein targets, which should result in persistent Tn antigen expression in the targeted cell types. Indeed, we observed Tn antigen expression on multiple glycoproteins in Far-Western blot format using the carbohydrate binding lectin *Vicia villosa* agglutinin (VVA), which binds well to α-linked GalNAc^44^ (**Fig. 4A**). Lectin blotting with VVA showed no detectible signal in wild-type brain, consistent with our prior studies showing a lack of significant levels of Tn antigen at baseline^18^ (**Fig. 4B**). Knock-out males from both *Cosmc Syn1-cKO* and *Cosmc Gfap-cKO* lines showed robust VVA binding in brain lysates across a range of molecular weights with distinct patterns, confirming inhibition of O-GalNAc extension that differed between lines.

**Figure 4.**
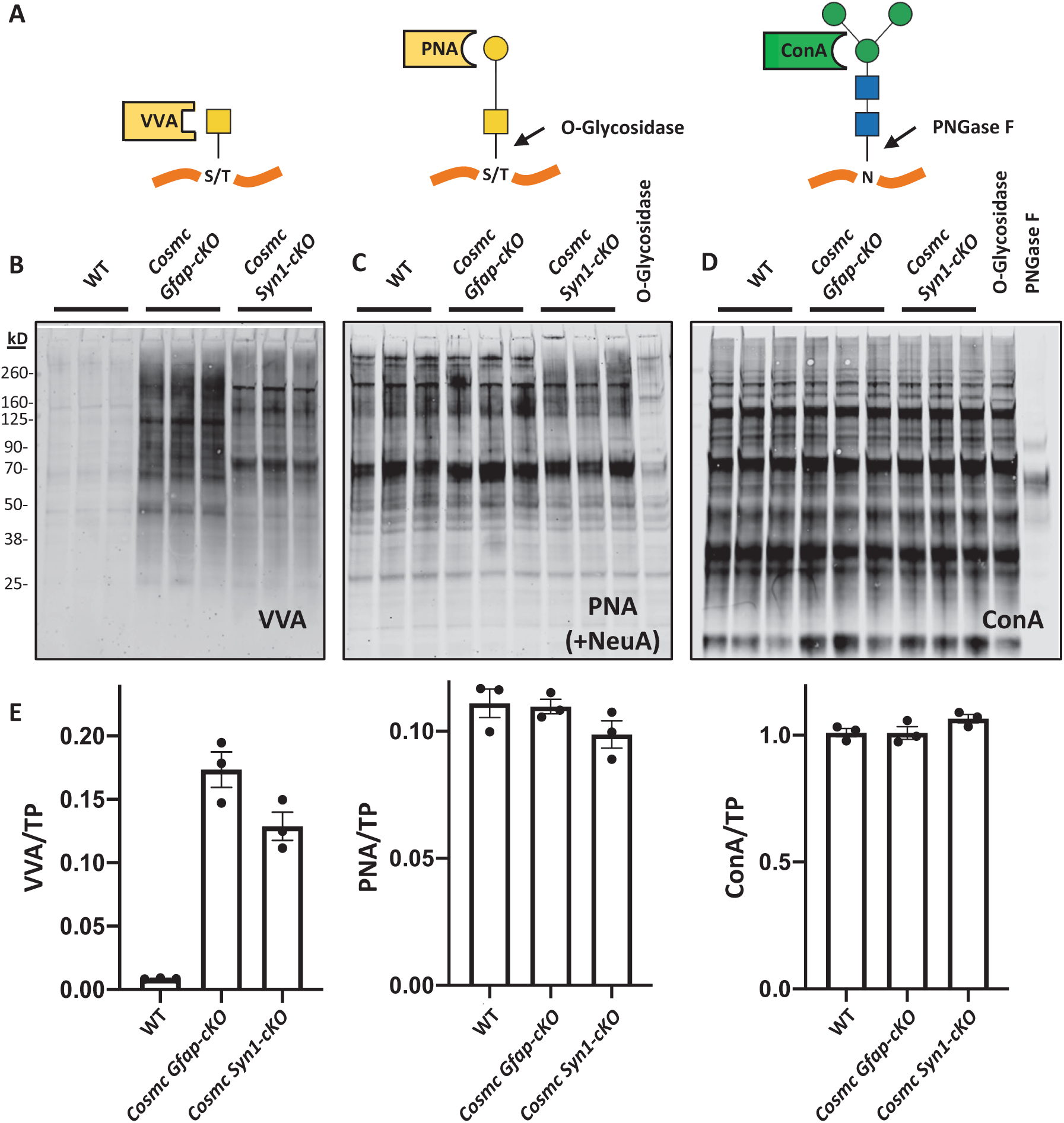
Genetic deletion of *Cosme* in *Syn1* and *Gfap* expressing cells had minimal effect on total O-GalNAc levels. **A)** Binding preference of multiple lectins and sites of glycosidase sensitivity: VVA binds to Tn-antigen; PNA binds T-antigen and is cleaved by O-glycosidase; ConA binds the core structure of N-glycans and is cleaved by PNGase F. **B)** VVA binding to endogenous Tn-antigen is absent in protein blots of wild type mouse cortex but generated in *Synl* and *Gfap Cosme-eKO* lines with distinct patterns, consistent with differential inhibition of O-GalNAc extension in these conditional knock-outs. **C)** Total levels of O-GalNAc glycans detected by PNA blotting after neuraminidase treatment showed no significant reduction in either cKO line, though the migration patterns of some protein bands differed compared to WT. **D)** ConA binding to N-glycans in cortex was unaffected in all mouse lines. Each lane contains cortical lysates from an individual mouse of the corresponding genotype. Experiment was replicated twice with similar results. **E)** Quantification of lectin binding from (B-D) normalized to total protein (TP) staining within the same lane of each blot. Data presented as the mean binding intensity +/- SEM, with data points for each individual mouse included. 15 µg protein lysate added per well.

Similar to lectin fluorescence, PNA binding to brain lysate on protein blots is dramatically increased by NeuA treatment and specific for O-glycans as the signal was reduced by O-glycosidase with or without NeuA (**Supplemental Fig. 6**), consistent with our mass spectrometry data showing that the majority of O-glycans in the mammalian brain are sialylated extensions of the T antigen^18^. Significant loss of extended O-glycans would be predicted to reduce PNA binding after removal of sialic acid with NeuA (**Fig. 2A**). Although some individual PNA positive bands were notably absent in the *Cosmc Syn1-cKO* deleted brains, including two large bands over 260 kD, no significant reduction of total PNA binding following NeuA treatment was observed by Western blotting in either *Syn1-* or *Gfap-Cosmc* deleted lines compared to wild-type mice (**Fig. 2C**). Binding of ConA, which is specific for high mannose, hybrid, and complex-type biantennary N-glycans^45,46^ and can be removed by treatment with Peptide:N-Glycosidase F (PNGase F)^47^, was not changed in the *Cosmc*-deleted lines, indicating that N-glycan biosynthesis was also not impacted by *Cosmc* deletion (**Fig. 4D**). When lectin binding was normalized to total protein from each lane, we observed consistent quantitative measures across multiple animals (**Fig. 4E**).

### PNA and VVA distribution in cerebellum showed distinct and inverse correlation in *Cosme eKO* lines

Though total PNA levels on lectin blotting were quantitatively unchanged in *Cosmc Syn1- and Gfap-cKO* lines, differences in PNA and VVA staining and migration patterns suggested O-GalNAcylation was altered distinctively in each model (**Fig. 2B**). Co-staining with common markers of astrocytes, neurons, and myelin (GFAP, NeuN, and MBP) in each *Cosmc cKO* line showed a preservation of macroscopic brain anatomy (**Supplemental Fig. 7**). Consistent with blotting studies, minimal VVA positivity was detected in the wild-type brain, and specificity was confirmed in the *Cosmc Syn1-cKO* line using competing levels of GalNAc to block all binding (**Supplemental Fig. 8**). VVA staining in *Cosmc Syn1-cKO* mice showed a speckled pattern with enrichment in white matter tracts of the cerebellum, while *Cosmc Gfap-cKO* mice showed diffuse staining with the least amount present in white matter tracts (**Fig. 5A**, **Supplemental Fig. 8**).

**Figure 5.**
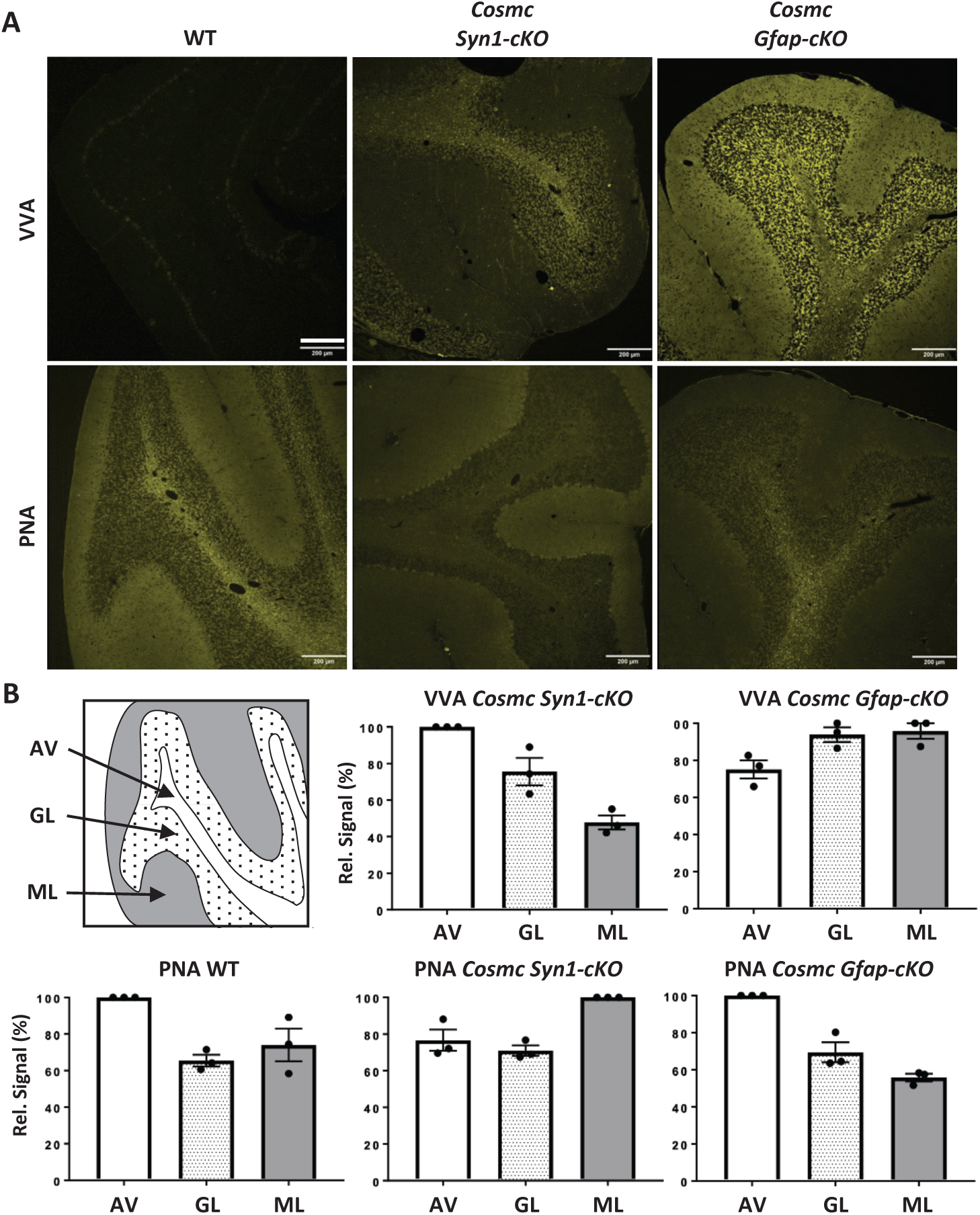
The distribution of VVA and PNA binding differs between cKO lines. **A)** VVA and PNA binding in adult mouse cerebellum of wild-type and *Cosme* deleted *Synl* and *Gfap* lines. Scale bar = 200 µm. **B)** Schematic illustrating different layers of the cerebellar including the arbor vitae (AV), granular layer (GL), and molecular layer (ML), and relative quantification of VVA and PNA binding between each layer from the three lines. Of note, no appreciable VVA binding is observed in wild-type brain. Data presented as the mean binding intensity +/- SEM, with individual data points from three independent mouse sections.

Compared to wild-type cerebellum, PNA staining in *Cosmc Syn1-cKO* mice was decreased in the arbor vitae, while in *Cosmc Gfap-cKO* mice it was reduced in the molecular layer (**Fig. 5B**). These findings suggest O-glycan synthesis differs between cell populations and an inverse spatial correlation between PNA and VVA binding is present following cell-specific *Cosmc* deletion.

### *Cosme* deletion in oligodendrocytes had minimal effect in brain

Given the enrichment of PNA signal in white matter and the lack of a severe phenotype and normal PNA levels in *Cosmc Syn1-cKO* and *Cosmc Gfap-cKO*, we considered whether our *Cosmc*-deletion strategy may have missed a critical brain cell type involved in O-GalNAc biology, specifically those of the oligodendrocyte lineage. Both *Gfap*- and *Syn1-Cre* expression lines have minimal expression in oligodendrocyte and polydendrocyte (**Supplemental Table 1**). To ensure inhibition of O-GalNAc extension in the oligodendrocyte lineage, we employed the established *Olig2-Cre* strain crossed in a similar strategy to floxed-*Cosmc* females^48^. Again, no overt phenotype was observed into adulthood, and normal birth ratios between male and female mice suggested that embryonic lethality was unlikely (**Supplemental Table 2**). On both lectin blot and fluorescent imaging, PNA and VVA staining in *Cosmc Olig2-cKO* cerebellum resembled that of wild-type mice (**Supplemental Fig. 9**). Taken together, these results suggest that global changes to the O-glycome and severe CDG phenotypes are not mirrored with single cell knock-out, and that more subtle perturbations and phenotypes may be associated with O-GalNAc biology in the brain.

### O-glycans are abundant on lecticans and display site-specific heterogeneity

Given the unique spatial distribution and migration patterns of PNA and VVA bound glycoproteins in wild-type and *Cosmc cKO* mice lines, we next sought to determine the identity of O-GalNAc carriers in the mammalian brain. Intact glycoproteins containing O-glycans from wild-type brain lysate after treatment with NeuA were affinity purified using PNA-bound agarose beads. After several washing steps, peptides were generated by trypsin digestion and analyzed by liquid chromatography-mass spectrometry (LC-MS/MS) (**Fig. 6A**). In total, 3,528 individual glycopeptides were detected. Analysis revealed 114 distinct glycan compositions, of which the most abundant species (H1N1A1 and H1N1) were consistent with T-synthase/Cosmc-mediated synthesis (**Fig. 6B**). While over 65% of NeuA treated and PNA precipitated O-glycans detected with LC-MS/MS lacked sialic acid, compared to over 90% of native O-glycans detected with MALDI^18^, nearly 35% contained at least 1 sialic acid (**Supplemental Fig. 10**). This result is suggestive of a partial desialylation reaction of the intact glycoproteins, as the most dominant species detected by MALDI in brain is the disialylated T-antigen, which was at only 7% in this study. The abundance of fucosylated O-glycans detected by LC-MS/MS and MALDI was similar (17% and 7%, respectively) (**Supplemental Fig. 10**).

**Figure 6.**
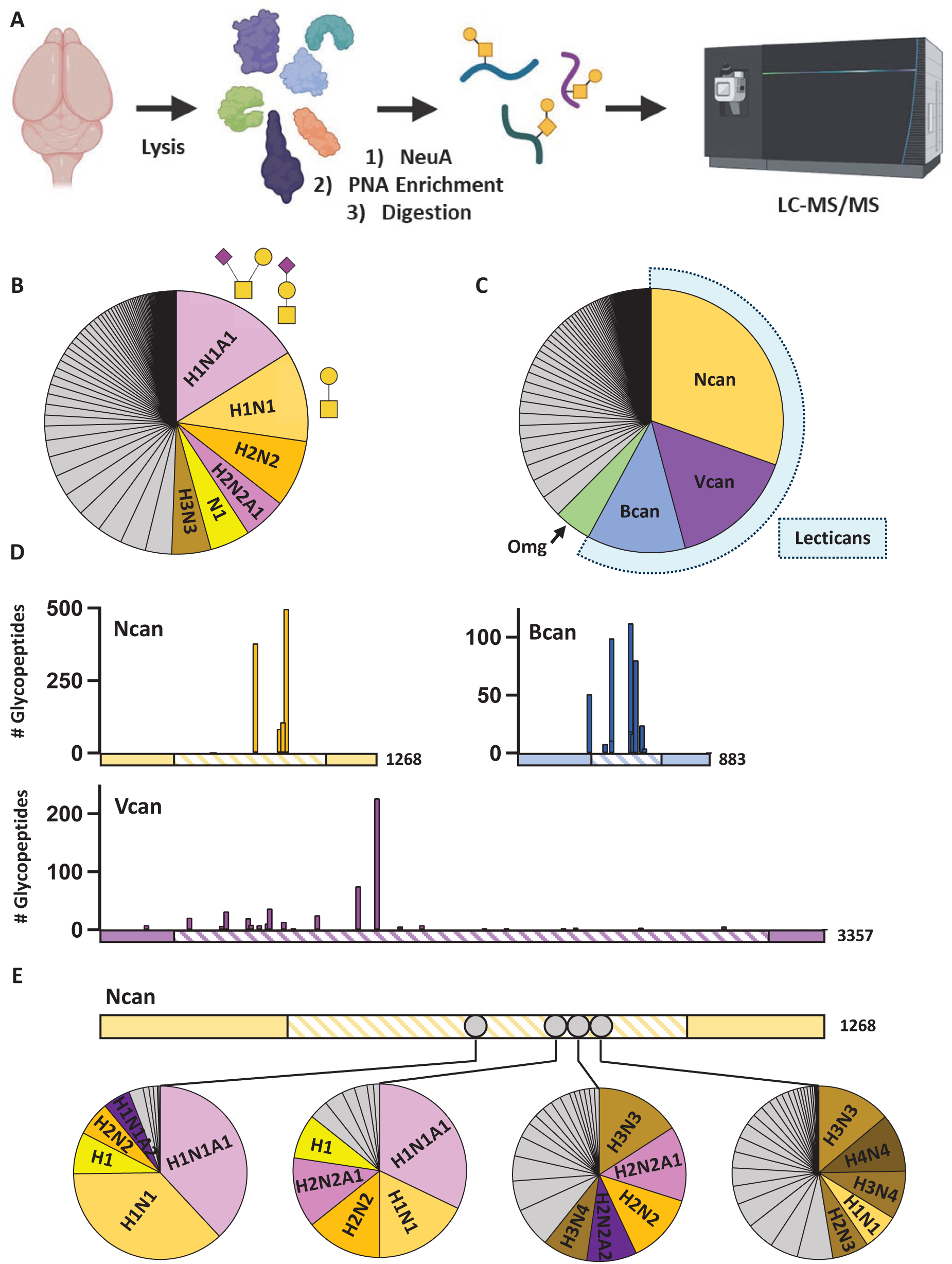
O-GalNAc are enriched on lecticans and display site-specific variability. **A)** Workflow to identify brain O-GalNAc glycans using PNA enrichment and LC-MS/MS. **B)** Of the 114 glycan compositions detected, the most abundant were consistent with core-1 GalNAc structures (H1N1A1, H1N1) common to the brain. **C)** Detected glycopeptides mapped on to 142 individual genes, with nearly 60% present on 3 proteins - Neurocan (Ncan), Brevican (Bcan), and Versican (Vcan) - members of the class of chondroitin-sulfate proteoglycans termed lecticans. **D)** Quantification of glycopeptide counts across the lecticans shows that most O-GalNAc sites were present in the non-homologous central domains (dashed lines), which also contain attachment sites for chondroitin sulfate chains. **E)** Analysis of composition from the 4 most abundant glycopeptides of Ncan demonstrated that each site contains distinct O-glycans. H = hexose; N = N-acetyl-hexosamine; A = NeuAc.

The glycopeptides mapped to 142 individual genes (**Fig. 6C**). Surprisingly, over half of the detected O-glycopeptides mapped to three proteins within the same family - Neurocan (Ncan), Brevican (Bcan), and Versican (Vcan). These proteins are members of the chondroitin sulfate proteoglycan family termed lecticans^49^, which are large extracellular scaffolding molecules enriched in structures including nodes of Ranvier^50^. Lecticans share multiple conserved domains including an N-terminal hyaluronic acid binding domain and C-terminal C-type lectin domain, and epidermal growth factor and complement regulatory protein like binding domains^49^. Between these conserved regions is a non-homologous central domain which contains attachment sites for glycosaminoglycan sites and varies in size between each lectican. Nearly all the O-glycans on lecticans were present in the central non-homologous domains (**Fig. 6D**).

Analysis of individual glycosites within Ncan which had more than 3 detected glycans showed that each displayed a distinct compositional profile (**Fig. 6E**). Many additional glycoproteins of considerable significance were modified by O-glycans, including oligodendrocyte myelin glycoprotein (Omg) and several cell adhesion and extracellular matrix molecules. Gene Ontology (GO) analysis of O-glycan carriers showed a significant enrichment for the synapse and processes including cell adhesion and development (**Supplemental Fig. 11**), suggesting a broad role of this modification in the brain.

### O-glycans are abundant at nodes of Ranvier and colocalize with Siglec-4 binding

Given their unique distribution in white matter and presence on extracellular matrices, we explored the relationship of O-glycans to the unmyelinated gaps of axons, termed nodes of Ranvier^51^. Co-labeling with PNA and an antibody to the paranodal marker Caspr^52^ demonstrated that the punctate signal of PNA localized directly within the node of Ranvier (**Fig. 7A**). Siglec-4, also known as Myelin-Associated Glycoprotein (MAG), is expressed by oligodendrocytes and involved in the maintenance and organization of the periaxonal region^53^ and in the development of the paranode region^54,55^. MAG is traditionally thought to bind sialic acid of gangliosides to mediate its function^56^, though cell-based binding assays showed that MAG binding was highest for the disialylated T-antigen of O-glycans^57^, which is the most abundant O-glycan in brain^18^. Binding of recombinant MAG to cerebellar brain tissue showed direct overlap with PNA, both enriched in white matter and with punctate staining (**Fig. 7B**).

**Figure 7.**
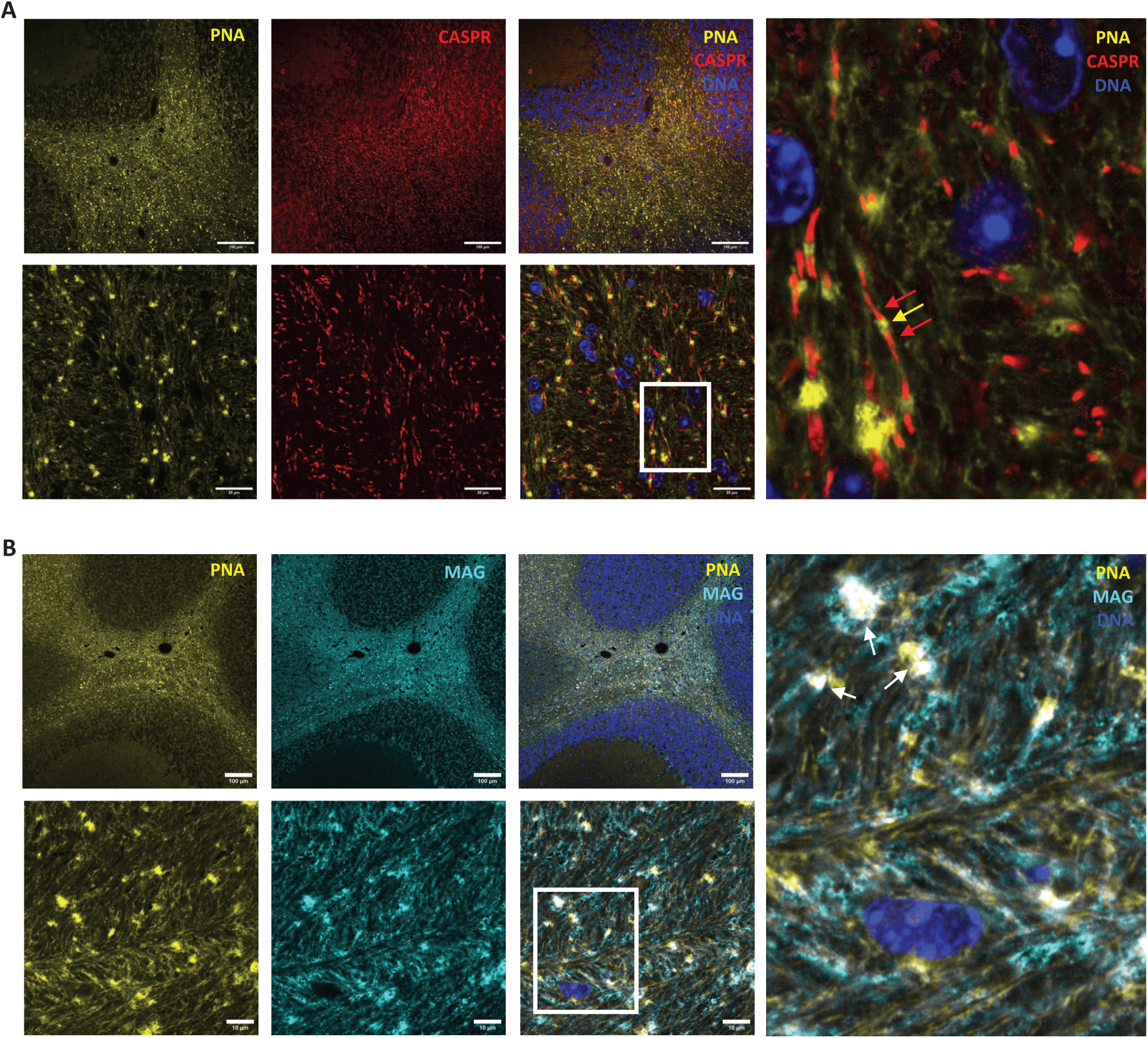
O-GalNAc glycans are abundant at nodes of Ranvier and colocalize with Siglec-4 (MAG) binding. **A)** PNA binding (yellow arrow) localizes between the antibody signal for the paranodal marker Caspr (red arrows). Top row scale bar = 100 µm, bottom row scale bar = 20 µm, with area enlarged in the right panel illustrated by a white box. **B)** PNA binding co-localizes with recombinant MAG (white arrows), a glycoprotein which binds sialylated glycans. Top row scale bar = 100 µm, bottom row scale bar = 10 µm, with area enlarged in the right panel illustrated by a white box.

### *Cosme* deletion in neurons reduces node of Ranvier length and Siglec-4 binding

Disruption of assembly mechanisms at nodes of Ranvier often lead to only a subtle effect, such as changes in length, while more profound phenotypes require disruption of multiple components^58^. We measured node of Ranvier length across wild-type and *Cosmc cKO* mice lines by measuring the distance between Caspr staining, as previously described^59^. Node of Ranvier length was significantly reduced in *Cosmc Syn1-cKO* mice by 20%, while unchanged in the other lines compared to wild-type (**Fig. 8A**). We next measured binding of recombinant MAG to brain tissue lysate immobilized on nitrocellulose and observed a reduced signal across multiple protein masses in *Cosmc Syn1-cKO* lysate (**Fig. 8B**). Quantification of MAG binding normalized to total protein staining showed a significant reduction in *Cosmc Syn1-cKO* lysate of ∼25% (**Fig. 8C**).

**Figure 8.**
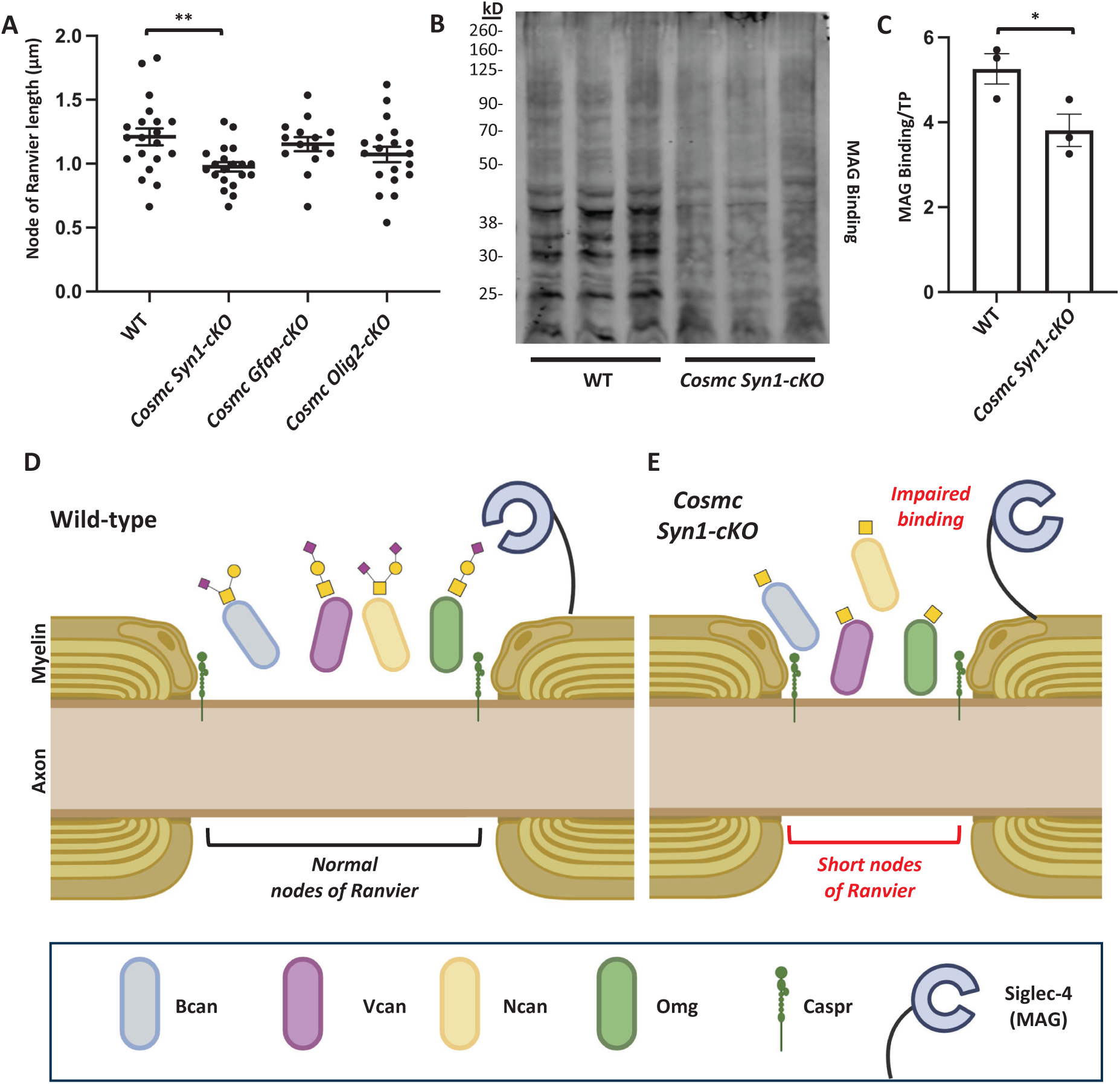
Inhibiting O-GalNAc synthesis in neurons shortens node of Ranvier length and reduces Siglec-4 (MAG) binding. **A)** Node of Ranvier length, measured as the distance between paranodal Caspr labeling, is significantly reduced in the *Cosme Synl-eKO* line compared to wild-type mice, but is unchanged in the other genotypes. Data shown from individual data points with the mean +/- SEM as horizontal lines. ** *p*-value = 0.003 using unpaired t-test. **B)** Binding of recombinant Siglec-4 (MAG) to immobilized brain protein was reduced in *Cosme Synl-eKO* lysate compared to wild-type. 15 µg of whole brain lysate was added per well, with technical triplicates from WT and *Cosme Synl-eKO* mice. **C)** Quantification of MAG binding from (B) normalized to total protein (TP) staining within the same lane. Data presented as the mean binding intensity +/- SEM, with data points for each sample included. * *p*-value = 0.049 using unpaired t-test. **D)** Model of the role of O-GalNAc glycans at the node of Ranvier. In wild-type mouse brain, O-GalNAc glycans are present on several lecticans (Ncan, Bcan, and Vcan) and Omg, and are a binding target of Siglec-4 (MAG). **E)** In *Cosme Synl-eKO* mouse brain, complex O-GalNAc synthesis is impaired, which leads to shortened length of the nodes of Ranvier and reduced binding of MAG.

## Discussion

Protein glycosylation in development and disease is a growing area of neuroscience research, but the total glycoproteome and its modifications are not well understood. Here we focused on O-GalNAc glycans (O-glycans) in brain glycoproteins, generating a comprehensive map of the cellular specificity, spatial distribution, and molecular identity of protein carriers of an abundant, yet perhaps least understood, O-glycosylation modification in the mammalian brain. We report that O-GalNAc glycans enrich in white matter tracts to the node of Ranvier and are present on several critical proteins including lecticans and others involved in brain development (**Fig. 8D**). Inhibition of O-GalNAc synthesis in cells expressing the *Syn1*, a synaptic protein with high levels in neurons, results in reduced length of nodes of Ranvier and decrease binding of the protein Siglec-4 (**Fig. 8E**). These findings suggest an important role for O-GalNAc glycans in the brain that has not previously been appreciated.

In previous studies, we predicted that the total abundance of O-glycans is somewhat less than N-glycans^7,18^, though genetic studies of both common and rare variants highlight the role of both pathways in disorders of the brain. The spatial distribution of O-glycans in the mouse brain differed starkly from that of N-glycans, as assessed by histological studies using lectins and knockout mice. We recently described the qualitative spatial distribution of N-glycan classes across brain regions after individually confirming their sensitivity and specificity using PNGase F cleavage and inhibition by blocking sugars^41^. Interestingly, N-glycans containing more terminal structures, such as fucose and bisected GlcNAc, showed enrichment in synapse dense regions including the molecular layer of the cerebellum^41^. These results were consistent with a recent findings using purified synaptosomes^60^. The different distribution of O-glycans compared to N-glycans may be partially accounted for by differential gene expression, given the slight enrichment of *T-Synthase* and *Cosmc* in oligodendrocyte and glial lineages from single cell databases (**Supplemental Table 1**), though we suspect other processes play a role in their unique spatial distribution, such as trafficking through the Golgi.

Global embryonic deletion of *Cosmc* in mice is lethal^22^, and we anticipated an overt phenotype resulting from genetic inhibition of the O-GalNAc pathway in the brain, such as poor growth or motor abnormalities, based on other studies in which this pathway has been blocked in other cell types and organs^22,32^. However, the change in node of Ranvier length alone without severe pathology is consistent with other studies involving assembly of this specialized structure, including cell-specific deletion of a critical transcription factors in oligodendrocytes^61^. Disruption of a single mechanism, such as the extracellular matrix, paranodal junction, or axonal cytoskeletal scaffold, results in mostly normal nodes, while disruption of any two mechanisms in combination results in motor dysfunction and early lethality^58^. Thus, a more severe phenotype of O-GalNAc inhibition may result from a combination of gene deletions or be exacerbated during disease states. A recent study using a novel lectin probe specific for the T antigen disaccharide Gal-GalNAc, showed localized binding to nodes of Ranvier, and showed dynamic binding during both brain development and demyelination^62^, though controls to demonstrate whether the epitope was on N- or O-linked glycans were not provided. Changes in node of Ranvier length are associated with differences in axonal conduction speed^59^, and may be altered in *Cosmc Syn1-cKO* mice.

The lack of a severe phenotype upon targeted deletion of *Cosmc* might also arise due to incomplete *Cre* expression/*Cosmc* deletion in cell lineages, though this seems unlikely given the generation of VVA positivity in each of the cKO lines. Germline recombination of *Cre*, which has been previously reported in other nervous system-specific expression systems^62^, as well as early embryonic lethality in our *Cosmc* KO lines, were both considered. However, given the normal birth ratios and multiple normal adult animals with blockage of O-glycan extension in the brain confirmed through both genotyping and VVA positivity, in addition to surviving mosaic female mice from the same litter, these potential confounds appear unlikely.

A functional role of O-glycans in the human brain is supported by our recent report of the *Cosmc*-CDG patients who exhibit neurologic abnormalities^34^. Our identification of this modification on numerous synaptic and cell adhesion molecules, in particular each of the brain-expressed lecticans, canonical carriers of proteoglycans, suggests a fundamental role in brain development and function. Our list of O-glycoproteins precipitated by PNA has considerable overlap with a recent study of brain O-glycoproteins measured using peptide fragments generated by a protease which cleaves immediately after O-GalNAc glycans^64^. The O-GalNAc modification may act competitively or synergistically with the well-established roles of different glycan modifications on the same protein.

Brevican is present at large nodes of Ranvier and regulates their assembly^65^, and previous work showed a dramatic migratory shift of certain brevican species on western blot following treatment with O-glycosidase and NeuA, consistent with the presence of multiple O-GalNAc glycans^66^. Of note, no cadherins or protocadherins were identified as carriers of O-GalNAc glycans in our study. At least two distinct O-mannosylation pathways exist for the cadherin superfamily^67^, both of which are present in the endoplasmic reticulum and may compete with Golgi O-GalNAc modification. O-glycome changes have been observed in human post-mortem brain samples affected by Parkinson’s and Lewy body disease, as well as mouse model harboring a common variant associated with schizophrenia risk, though both of these analyses included O-GalNAc and O-Mannose glycans together released through similar protocols^7,17^.

The macrophage galactose-type lectin (MGL), an endogenous C-type lectin expressed by macrophages, can bind to the Tn antigen^68^ and is involved in the recovery phase following demyelination in a model of multiple sclerosis^69^. Siglec-4, which is expressed by oligodendrocytes and is involved in axon and nodal development, binds sialylated T antigen in cell based assays^57^. We identified colocalization of Siglec-4 binding with O-glycans, and reduced Siglec-4 binding to *Cosmc Syn1-cKO* brain lysate. These findings further support a role of O-GalNAc synthesis and elongation in develop and disease of the brain. It has been observed that dynamic changes of PNA binding arise following acute depolarizing stimuli of brain slices^70^. Although it was not commented on in that study, PNA binding in the hippocampus and corpus callosum shows a similar distribution pattern with enrichment in white matter tracts. Following depolarization with high potassium conditions, PNA binding increased dramatically. However, at this time, without additional controls showing the specificity of PNA binding such as sensitivity of the signal to O-glycosidase or competition from galactose, we are cautious in drawing conclusions about PNA binding in this setting.

Our experimental design leveraged *Cosmc* deletion as a bottleneck to inhibit the O-GalNAc pathway in the brain, and as *Cosmc* is expressed on the X chromosome, this approach places primary focus on male cKO mice as a model. Previous studies of *Cosmc* deletion have leveraged mosaic females in the setting where global deletion in males was lethal or the phenotype was too severe, though this was not the case in our brain *Cosmc* cKO lines as both hemizygous knockout males and mosaic females were grossly normal. Further, recent studies reveal that *COSMC*-CDG is an X-linked condition primarily affecting males^34^. Thus, while a limitation of our study is focusing on male mice to leverage the genetic vulnerability and simplicity of O-GalNAc inhibition, future studies targeting O-glycans in females are warranted. Future studies of O-glycans in white matter will advance our understanding of their involvement in development of diseases of the brain, which are clearly implicated by the growing list of genetic conditions with neuropsychiatric phenotypes associated with enzymes important to O-GalNAc glycan biosynthesis.

## Methods

All mice were maintained in standard housing conditions, provided normal chow and water *ad lib*, and maintained on 12-hour day/light cycle. All mice were housed and maintained in accordance with the guidelines established by the Animal Care and Use Committee at Beth Israel Deaconess Medical Center under the approved protocol #02-2022.

Homozygous floxed female Cosmc mice on C57/BL6 background generated in a previous study^22^ were crossed with male Cre-driver lines obtained from Jackson laboratories including *Syn1-Cre* (strain:003966), *Gfap-Cre* (strain:024098), and *Olig2-Cre* (strain:025567), all on C57/BL6 backgrounds. Twelve-week-old male mice were used in this study unless otherwise indicated.

Mouse genotyping was performed using the following primers as described in prior studies^22^ and per recommendation of Jackson Laboratories. *Cosmc*: Forward 5’-GCA ACA CAA AGA AAC CCT GGG - 3’, Reverse 5’ - TCG TCT TTG TTA GGG GCT TGC -3’; WT allele is 774 bp and floxed Cosmc allele is 808 bp. *Syn1*: Forward 5’ - CTC AGC GCT GCC TCA GTC T - 3’, Reverse 5’ - GCA TCG ACC GGT AAT GCA - 3’; Transgene amplifies 300 bp product. *Gfap*: Forward 5’ - TCC ATA AAG GCC CTG ACA TC - 3’, Reverse 5’ - TGC GAA CCT CAT CAC TCG T - 3’; Transgene amplifies 400 bp product.

*Brain dissection*, *tissue lysis*, and *lectin-blotting* were performed as previously described^18^, with the following modifications. For lectin blotting with PNA, 50 µg of protein lysate was treated with 5 µL NeuA (a2-3,6,8,9 Neuraminidase A, P0722L, NEB) or 5 µL of O-Glycosidase (P0733L, NEB) where indicated prior to denaturation/loading on SDS-PAGE gels according to manufacturer protocols. Membranes were incubated with 2 µg/mL biotinylated lectins from Vector Labs (VVA B-1235-2, PNA B-1075-5) in conditions as previously described, imaged using a LiCOR Odyssey CLx Imaging System and analyzed using LiCOR Image Studio Software. A comprehensive characterization of biotinylated lectin binding specificity by glycan microarray can be found on the website for the National Center for Functional Glycomics.

### Histology/immunofluorescence

Mice at 12 weeks of age were euthanized with CO_2_ gas in accordance with AVMA guidelines and trans-cardially perfused with ice-cold PBS containing heparin at 10,000 units/L (Sigma #3149) for 2 minutes at 6 ml/minute, followed by ice-cold 4% PFA in 0.1 M PBS, pH 7.4 for 5 minutes at 5 ml/minute. Whole brains were then removed from the skull and postfixed in 4% PFA overnight and transferred to PBS until processing. Fixed brain was paraffin embedded and cut in 3 µm coronal sections with a microtome at the BIDMC Pathology Core Facility. Slides were then deparaffinized using a standard xylene/ethanol gradient protocol and placed in antigen retrieval solution (0.1 M citric acid and 0.1 M sodium citrate, pH=6), incubated in pressure cooker at boiling temperature for 3 min, and stored at 4°C in TBS. Tissue sections were circled by hydrophobic PAP PEN and slides were treated with denaturing buffer for 5 min at 95°C. Sections were washed with TBS (Trizma 20 mM, NaCl 100 mM, CaCl_2_ 1 mM, MgCl_2_ 1 mM, pH 7.2) 3 times followed by 3 washes with TBS-tween 0.05% (TBS-T). Sections were blocked with 3% BSA in TBS for 1 h at room temperature (RT). Tissues were washed again three times with TBS and incubated with 25 µg/mL of lectins (Vector Labs: ConA FL-1001, GNL B-1245, AAL FL-1391, PHA-E FL-1121, RCA-I B-1085, PNA FL-1071, VVA FL-1231) or a-NeuN (Abcam, ab177487, 0.876 mg/mL), a-GFAP (Sigma, HPA056030, 0.05 mg/mL), a-Caspr (Abcam, Ab34151, 1 mg/mL), a-MBP AF594-conjugated (Biolegend, 850905, 0.5 mg/mL) diluted at 1/100 in TBS-T or with MAG (Bio-Techne, 8580-MG, 200 µg/mL) diluted at 1/10 in TBS-T for 1 h at RT in darkness with gentle shaking. For control glycosidase experiments, slides were pretreated with 2,500 U of PNGase F, 200 U of NeuA, or 160,000 U of O-Glycosidase in a final volume of 80 µL at 37°C overnight prior to incubation with lectins. For control carbohydrate/haptenic sugar blocking experiments, lectins were incubated for 30 min with the indicated sugar before incubation on the slide for 1 hour at RT. After the incubation with lectins, sections were washed with TBS-T 3 times. Most lectins described above were directly conjugated to FITC - however, for biotinylated RCA-I and GNL, streptavidin-488 (Invitrogen, S11223) was used as secondary antibody at 10 µg/mL and incubated in TBS-T for 1 hour at 25°C prior to imaging. For immunofluorescence with a-NeuN and a-GFAP, a donkey anti-rabbit 647 secondary (Southern biotech, 6440-31, 1 mg/mL) was used diluted at 1/200 in TBS-T. For MAG and a-Caspr, an anti-mouse 488 (Invitrogen, A28175, 1 mg/mL) and anti-rabbit (Invitrogen, A10042, 2 mg/mL) were respectively diluted in 1/100 and 1/200 in TBS-T. Sections were washed 2 times with TBS-T, one time with TBS and counterstained with Hoechst 33342 diluted at 1/1000 in TBS for 10 min at RT in darkness. Sections were then washed 3 times with TBS, incubated with 0.1% Sudan Black B (Sigma, 199664) in 70% ethanol for 5 min to reduce autofluorescence coming from lipofuscin, washed again 2 times with TBS and mounted with a glass coverslip using Prolong Gold Antifade Mountant (ThermoFisher, P36930). After slides cured overnight, image acquisition was performed on Zeiss LSM 880 Upright Confocal System at the BIDMC Confocal Microscopy Core for high magnification and a VS120 Slidescanner from Olympus for low magnification. Laser intensity was unchanged between the experimental conditions and controls, for example +/- PNGase F. Images analysis was then performed using ImageJ (v1.53t) or QuPath (v0.3.2) software.

### Node of Ranvier length measurement

Node of Ranvier length measurements were performed as previously described^59^. Briefly, confocal images of cerebellum were analyzed using ImageJ software. Node diameters were measured using the line tool in ImageJ over Caspr staining as the distance between the half maximum intensity for each staining.

### Gene set and single cell expression analysis

Brain Single cell expression data for *T-synthase* (*C1galt1*), *Cosmc* (*C1galt1c1*), *Syn1*, and *Gfap* were downloaded from the DropViz database^36^ on December 10^th^, 2022.

### Glycoproteomics

Mouse brains were lyzed with a Dounce homogenizer in a lysis buffer containing 50 mM HEPES pH=7.8, 0.5% sodium deoxycholate, and cOmplete Protease Inhibitor Cocktail EDTA-free (Roche) at 4°C. The lysate was sequentially sonicated with a Brinkmann homogenizer three times, 20 A, 10 s, with 30 s rest on ice between rounds. Protein concentration was measured with BCA assay (Pierce). The lysate was then treated with 20k units of a2-3,6,8 neuraminidase (New England BioLabs) overnight at 37°C with end-over-end rotation. Peanut agglutinin (PNA) bound to agarose (Vector) was washed with water three times and transferred to the lysate. Calcium chloride and manganese chloride were also added to the enrichment to the final concentrations of 0.1 mM and 0.01 mM, respectively. The enrichment was performed at room temperature overnight with end-over-end rotation. The beads were washed three times with water. Enriched glycoproteins were eluted by incubation with a solution containing 200 mM galactose in 10 mM HEPES pH=7.8 for 30 min with end-over-end rotation. The eluted glycoproteins were purified with the methanol/chloroform precipitation. The pellet was dried and resuspended in a digestion buffer containing 1.6 M urea, 50 mM HEPES, pH=7.8, and 5% acetonitrile. The glycoproteins were digested with sequencing-grade modified trypsin (Promega) at 37°C with shaking overnight. The glycopeptides were desalted with a C18 Sep-Pak cartridge (Waters) and dried with lyophilization. Glycopeptides were fractionated with high-pH reversed-phase chromatography using a Shimadzu 20A HPLC system. Peptides were separated on an XBridge C18 3.5 µm, 4.6x250 mm column (Waters). Buffer A was 10 mM ammonium formate pH=10 in water and buffer B was 10 mM ammonium formate pH=10 in 90% ACN. The peptides were fractionated and collected during a 50-minute gradient of 5% to 35% buffer B and the flow rate of 0.7 mL/minute. The peptides were collected every 2 minutes, consolidated into 10 fractions, and dried with lyophilization. The peptides were desalted with a C18 Sep-Pak cartridge and dried with lyophilization.

Peptide sequencing was performed with an Orbitrap Fusion Lumos Tribrid mass spectrometer (Thermo). The glycopeptides were dissolved in a solution containing 0.1% formic acid (FA) in water and loaded onto a C18 precolumn (C18 PepMap 100, 300 µm x 5 mm, 5 µm, 100 A, Thermo) with 15 µL/min solvent A (0.1% FA in water) for 3 min, and separated on a C18 analytical column (PicoFrit 75 µm ID x 150 mm, 3 µm, New Objective) using a gradient of 2-40% solvent B (80% ACN and 0.1% FA in water) over 95 min, followed by a gradient of 40-90% over 3 min. The ion source voltage was 2100 V. The ion transfer tube temperature was 275°C. Higher-energy dissociation product ions-triggered electron-transfer/higher-energy collision dissociation (HCD-pd-EThcD MS2) was used for glycopeptide identification. The mass spectrometer was operated in a data-dependent acquisition mode where 15 most intense ions from MS1 are sequenced in MS2 with HCD, and those with specified oxonium ions in MS2 are also sequenced with EThcD at MS2 level. MS1 was performed in the Orbitrap at the resolution of 120,000. Depending on the experiment, the MS1 scan range was set to m/z 350-2000 or m/z 800-2000. Normalized AGC target was set 100% with the maximum ion injection time of 50 ms. The quadrupole was used to isolate precursor ions with the isolation window of 1.6 m/z. Dynamic exclusion was employed for 10 s with the mass tolerance of 10 ppm after 1 time. Stepped HCD with collision energies of 20%, 30%, and 40% was used to fragment the peptides in HCD MS2. MS2 detection was performed in the Orbitrap with the resolution of 30,000 and the first m/z of 100. The normalized AGC target was set to 100% with the maximum injection time of 250 ms. Targeted mass trigger was used after HCD MS2 where at least two ions with m/z=204.0865, 168.0654, 186.0760, 274.0921, 292.1027, 366.1395, 126.0550, 138.0549, or 144.0655, with mass tolerance of 25 ppm and are one of the 20 most intense ions triggered another MS2 with EThcD. Calibrated charge-dependent ETD was enabled with ETD supplemental energy of 25%. Ions were detected in the Orbitrap with a 30,000 resolution in the high mass range mode with the scan range from m/z=120-4000. The normalized AGC target was 200% and the maximum ion injection time was 200 ms. Raw files were searched using pGlyco3^71^ against the mouse proteome database (downloaded 06/09/2021, reviewed, containing both canonical and non-canonical sequences, 25368 entries) from UniProt. Trypsin was set as an enzyme with full specificity. Maximum number of missed cleavages was 2. Oxidation of methionine (+15.9949 Da) was set as a variable protein modification. Maximum variable modification was 2. Peptide length was from 6 to 40. The glycan database used was the default “Multi-Site-O-Glycan”. Precursor ion mass tolerance was 10 ppm and fragment ion mass tolerance was 20 ppm. The glycopeptide false discovery rate (FDR) was 1%. Identified glycopeptides were inspected and all were used for the analysis. Raw files are available on ProteomeXchange with accession number PXD041631.

### Gene set and single cell expression analysis

Brain Single cell expression data from the frontal cortex containing 14 distinct cell clusters was downloaded from the DropViz database^36^ on December 10^th^, 2022. Data for *T-synthase* (*C1galt1*), *Cosmc* (*C1galt1c1*), *Syn1*, *Gfap* and the 42 PNA-enriched glycoproteins were included, and exponentially natural log transformed to approximate transcripts per 100,000 in each cell cluster. Gene set enrichment analysis and figures were generated using the FUMA GWAS GENE2FUNC online tool^72^ on January 9, 2024, with adjusted P-value threshold set at < 0.05.

### Statistical Analysis

Statistical analysis of lectin blots and staining was performed using GraphPad Prism v8.4 and Microsoft Excel Version 16.72 using significance threshold set of < 0.05. Graphics were created using Microsoft Excel and PowerPoint Version 16.72, BioRender.com, and GraphPad Prism v8.4.

### Data Availability

The glycoproteomics datasets generated during and/or analysed during the current study are available in the ProteomeXchange Consortium via the PRIDE partner repository with the dataset identifier PXD041631. All additional datasets are available from the corresponding author on reasonable request.

## Acknowledgements

The authors would like to thank the Neurobiology Imaging Facility (NIF) at Harvard Medical School, Boston, MA (HMS/BCH CENTER FOR NEUROSCIENCE RESEARCH #NS072030), the Confocal Imaging Core and Histology Core at the Beth Israel Deaconess Medical Center for resources, technology, and expertise. RGM is supported by 1K08MH128712 and a 2021 NARSAD Young Investigator Grant from the Brain & Behavior Research Foundation. MN received a scholarship supplement from the Philippe Foundation. This research was supported by NIH Grant R24GM137763, awarded to RDC.

## Author contributions

NM, RDC, and RGM conceptualized the project and wrote the manuscript. NM and RGM carried out the majority of experiments and analysis. SS performed glycoproteomics analysis. RDC and RGM supervised the project. All authors contributed feedback and edits to the manuscript.

## References

1. Reily, C., Stewart, T.J., Renfrow, M.B., and Novak, J. (2019). Glycosylation in health and disease. Nat. Rev. Nephrol. 15, 346–366. 10.1038/s41581-019-0129-4.

2. Iqbal, S., Ghanimi Fard, M., Everest-Dass, A., Packer, N.H., and Parker, L.M. (2019). Understanding cellular glycan surfaces in the central nervous system. Biochem. Soc. Trans. 47, 89–100. 10.1042/BST20180330.

3. Hanus, C., Geptin, H., Tushev, G., Garg, S., Alvarez-Castelao, B., Sambandan, S., Kochen, L., Hafner, A.-S., Langer, J.D., and Schuman, E.M. (2016). Unconventional secretory processing diversifies neuronal ion channel properties. eLife 5, e20609. 10.7554/eLife.20609.

4. Schwartz, N.B., and Domowicz, M.S. (2018). Proteoglycans in brain development and pathogenesis. FEBS Lett. 592, 3791–3805. 10.1002/1873-3468.13026.

5. Freeze, H.H., Eklund, E.A., Ng, B.G., and Patterson, M.C. (2015). Neurological aspects of human glycosylation disorders. Annu. Rev. Neurosci. 38, 105–125. 10.1146/annurev-neuro-071714-034019.

6. Joshi, H.J., Hansen, L., Narimatsu, Y., Freeze, H.H., Henrissat, B., Bennett, E., Wandall, H.H., Clausen, H., and Schjoldager, K.T. (2018). Glycosyltransferase genes that cause monogenic congenital disorders of glycosylation are distinct from glycosyltransferase genes associated with complex diseases. Glycobiology 28, 284–294. 10.1093/glycob/cwy015.

7. Mealer, R.G., Williams, S.E., Noel, M., Yang, B., D’Souza, A.K., Nakata, T., Graham, D.B., Creasey, E.A., Cetinbas, M., Sadreyev, R.I., et al. (2022). The schizophrenia-associated variant in SLC39A8 alters protein glycosylation in the mouse brain. Mol. Psychiatry. 10.1038/s41380-022-01490-1.

8. Mealer, R.G., Williams, S.E., Daly, M.J., Scolnick, E.M., Cummings, R.D., and Smoller, J.W. (2020). Glycobiology and schizophrenia: a biological hypothesis emerging from genomic research. Mol. Psychiatry, 1–11. 10.1038/s41380-020-0753-1.

9. Suttapitugsakul, S., Stavenhagen, K., Donskaya, S., Bennett, D.A., Mealer, R.G., Seyfried, N.T., and Cummings, R.D. (2022). Glycoproteomics Landscape of Asymptomatic and Symptomatic Human Alzheimer’s Disease Brain. Mol. Cell. Proteomics MCP 21, 100433. 10.1016/j.mcpro.2022.100433.

10. Williams, S.E., Mealer, R.G., Scolnick, E.M., Smoller, J.W., and Cummings, R.D. (2020). Aberrant glycosylation in schizophrenia: a review of 25 years of post-mortem brain studies. Mol. Psychiatry, 1–10. 10.1038/s41380-020-0761-1.

11. Sipione, S., Monyror, J., Galleguillos, D., Steinberg, N., and Kadam, V. (2020). Gangliosides in the Brain: Physiology, Pathophysiology and Therapeutic Applications. Front. Neurosci. 14, 572965. 10.3389/fnins.2020.572965.

12. Conroy, L.R., Hawkinson, T.R., Young, L.E.A., Gentry, M.S., and Sun, R.C. (2021). Emerging roles of N-linked glycosylation in brain physiology and disorders. Trends Endocrinol. Metab. TEM 32, 980–993. 10.1016/j.tem.2021.09.006.

13. Klaric, T.S., and Lauc, G. (2022). The dynamic brain N-glycome. Glycoconj. J. 39, 443–471. 10.1007/s10719-022-10055-x.

14. Joshi, H.J., Narimatsu, Y., Schjoldager, K.T., Tytgat, H.L.P., Aebi, M., Clausen, H., and Halim, A. (2018). SnapShot: O-Glycosylation Pathways across Kingdoms. Cell 172, 632–632.e2. 10.1016/j.cell.2018.01.016.

15. Sheikh, M.O., Halmo, S.M., and Wells, L. (2017). Recent advancements in understanding mammalian O-mannosylation. Glycobiology 27, 806–819. 10.1093/glycob/cwx062.

16. Lee, B.E., Suh, P.-G., and Kim, J.-I. (2021). O-GlcNAcylation in health and neurodegenerative diseases. Exp. Mol. Med. 53, 1674–1682. 10.1038/s12276-021-00709-5.

17. Wilkinson, H., Thomsson, K.A., Rebelo, A.L., Hilliard, M., Pandit, A., Rudd, P.M., Karlsson, N.G., and Saldova, R. (2021). The O-Glycome of Human Nigrostriatal Tissue and Its Alteration in Parkinson’s Disease. J. Proteome Res. 20, 3913–3924. 10.1021/acs.jproteome.1c00219.

18. Williams, S.E., Noel, M., Lehoux, S., Cetinbas, M., Xavier, R.J., Sadreyev, R.I., Scolnick, E.M., Smoller, J.W., Cummings, R.D., and Mealer, R.G. (2022). Mammalian brain glycoproteins exhibit diminished glycan complexity compared to other tissues. Nat. Commun. 13, 275. 10.1038/s41467-021-27781-9.

19. Brockhausen, I., Wandall, H.H., Hagen, K.G.T., and Stanley, P. (2022). O-GalNAc Glycans. In Essentials of Glycobiology, A. Varki, R. D. Cummings, J. D. Esko, P. Stanley, G. W. Hart, M. Aebi, D. Mohnen, T. Kinoshita, N. H. Packer, J. H. Prestegard, et al., eds. (Cold Spring Harbor Laboratory Press).

20. Varki, A., Cummings, R.D., Aebi, M., Packer, N.H., Seeberger, P.H., Esko, J.D., Stanley, P., Hart, G., Darvill, A., Kinoshita, T., et al. (2015). Symbol Nomenclature for Graphical Representations of Glycans. Glycobiology 25, 1323–1324. 10.1093/glycob/cwv091.

21. Neelamegham, S., Aoki-Kinoshita, K., Bolton, E., Frank, M., Lisacek, F., Lutteke, T., O’Boyle, N., Packer, N.H., Stanley, P., Toukach, P., et al. (2019). Updates to the Symbol Nomenclature for Glycans guidelines. Glycobiology 29, 620–624. 10.1093/glycob/cwz045.

22. Wang, Y., Ju, T., Ding, X., Xia, B., Wang, W., Xia, L., He, M., and Cummings, R.D. (2010). Cosmc is an essential chaperone for correct protein O-glycosylation. Proc. Natl. Acad. Sci. 107, 9228–9233. 10.1073/pnas.0914004107.

23. Xia, L., and McEver, R.P. (2006). Targeted disruption of the gene encoding core 1 beta1-3-galactosyltransferase (T-synthase) causes embryonic lethality and defective angiogenesis in mice. Methods Enzymol. 416, 314–331. 10.1016/S0076-6879(06)16021-8.

24. Kudelka, M.R., Hinrichs, B.H., Darby, T., Moreno, C.S., Nishio, H., Cutler, C.E., Wang, J., Wu, H., Zeng, J., Wang, Y., et al. (2016). Cosmc is an X-linked inflammatory bowel disease risk gene that spatially regulates gut microbiota and contributes to sex-specific risk. Proc. Natl. Acad. Sci. U. S. A. 113, 14787–14792. 10.1073/pnas.1612158114.

25. Wang, Y., Jobe, S.M., Ding, X., Choo, H., Archer, D.R., Mi, R., Ju, T., and Cummings, R.D. (2012). Platelet biogenesis and functions require correct protein O-glycosylation. Proc. Natl. Acad. Sci. U. S. A. 109, 16143–16148. 10.1073/pnas.1208253109.

26. Fu, J., Gerhardt, H., McDaniel, J.M., Xia, B., Liu, X., Ivanciu, L., Ny, A., Hermans, K., Silasi-Mansat, R., McGee, S., et al. (2008). Endothelial cell O-glycan deficiency causes blood/lymphatic misconnections and consequent fatty liver disease in mice. J. Clin. Invest. 118, 3725–3737. 10.1172/JCI36077.

27. Stotter, B.R., Talbot, B.E., Capen, D.E., Artelt, N., Zeng, J., Matsumoto, Y., Endlich, N., Cummings, R.D., and Schlondorff, J.S. (2020). Cosmc-dependent mucin-type O-linked glycosylation is essential for podocyte function. Am. J. Physiol. Renal Physiol. 318, F518–F530. 10.1152/ajprenal.00399.2019.

28. Cutler, C.E., Jones, M.B., Cutler, A.A., Mener, A., Arthur, C.M., Stowell, S.R., and Cummings, R.D. (2019). Cosmc is required for T cell persistence in the periphery. Glycobiology. 10.1093/glycob/cwz054.

29. Zeng, J., Aryal, R.P., Stavenhagen, K., Luo, C., Liu, R., Wang, X., Chen, J., Li, H., Matsumoto, Y., Wang, Y., et al. (2021). Cosmc deficiency causes spontaneous autoimmunity by breaking B cell tolerance. Sci. Adv. 7, eabg9118. 10.1126/sciadv.abg9118.

30. Zeng, J., Eljalby, M., Aryal, R.P., Lehoux, S., Stavenhagen, K., Kudelka, M.R., Wang, Y., Wang, J., Ju, T., von Andrian, U.H., et al. (2020). Cosmc controls B cell homing. Nat. Commun. 11, 3990. 10.1038/s41467-020-17765-6.

31. Trubetskoy, V., Pardifias, A.F., Qi, T., Panagiotaropoulou, G., Awasthi, S., Bigdeli, T.B., Bryois, J., Chen, C.-Y., Dennison, C.A., Hall, L.S., et al. (2022). Mapping genomic loci implicates genes and synaptic biology in schizophrenia. Nature 604, 502–508. 10.1038/s41586-022-04434-5.

32. Zilmer, M., Edmondson, A.C., Khetarpal, S.A., Alesi, V., Zaki, M.S., Rostasy, K., Madsen, C.G., Lepri, F.R., Sinibaldi, L., Cusmai, R., et al. (2020). Novel congenital disorder of O-linked glycosylation caused by GALNT2 loss of function. Brain 143, 1114–1126. 10.1093/brain/awaa063.

33. Gollamudi, S., Lekhraj, R., Lalezari, S., and Lalezari, P. (2020). COSMC mutations reduce T-synthase activity in advanced Alzheimer’s disease. Alzheimers Dement. N. Y. N 6, e12040. 10.1002/trc2.12040.

34. Erger, F., Aryal, R.P., Reusch, B., Matsumoto, Y., Meyer, R., Zeng, J., Knopp, C., Noel, M., Muerner, L., Wenzel, A., et al. (2023). Germline C1GALT1C1 mutation causes a multisystem chaperonopathy. Proc. Natl. Acad. Sci. U. S. A. 120, e2211087120. 10.1073/pnas.2211087120.

35. Itoh, K., and Nishihara, S. (2021). Mucin-Type O-Glycosylation in the Drosophila Nervous System. Front. Neuroanat. 15, 767126. 10.3389/fnana.2021.767126.

36. Saunders, A., Macosko, E.Z., Wysoker, A., Goldman, M., Krienen, F.M., de Rivera, H., Bien, E., Baum, M., Bortolin, L., Wang, S., et al. (2018). Molecular Diversity and Specializations among the Cells of the Adult Mouse Brain. Cell 174, 1015–1030.e16. 10.1016/j.cell.2018.07.028.

37. Lotan, R., Skutelsky, E., Danon, D., and Sharon, N. (1975). The purification, composition, and specificity of the anti-T lectin from peanut (Arachis hypogaea). J. Biol. Chem. 250, 8518–8523.

38. Cummings, R.D., Etzler, M., Hahn, M.G., Darvill, A., Godula, K., Woods, R.J., and Mahal, L.K. (2022). Glycan-Recognizing Probes as Tools. In Essentials of Glycobiology, A. Varki, R. D. Cummings, J. D. Esko, P. Stanley, G. W. Hart, M. Aebi, D. Mohnen, T. Kinoshita, N. H. Packer, J. H. Prestegard, et al., eds. (Cold Spring Harbor Laboratory Press).

39. Fujita, K., Oura, F., Nagamine, N., Katayama, T., Hiratake, J., Sakata, K., Kumagai, H., and Yamamoto, K. (2005). Identification and molecular cloning of a novel glycoside hydrolase family of core 1 type O-glycan-specific endo-alpha-N-acetylgalactosaminidase from Bifidobacterium longum. J. Biol. Chem. 280, 37415–37422. 10.1074/jbc.M506874200.

40. Bojar, D., Meche, L., Meng, G., Eng, W., Smith, D.F., Cummings, R.D., and Mahal, L.K. (2022). A Useful Guide to Lectin Binding: Machine-Learning Directed Annotation of 57 Unique Lectin Specificities. ACS Chem. Biol. 17, 2993–3012. 10.1021/acschembio.1c00689.

41. Noel, M., Cummings, R.D., and Mealer, R.G. (2023). N-glycans show distinct spatial distribution in mouse brain. Glycobiology, cwad077. 10.1093/glycob/cwad077.

42. Zhu, Y., Romero, M.I., Ghosh, P., Ye, Z., Charnay, P., Rushing, E.J., Marth, J.D., and Parada, L.F. (2001). Ablation of NF1 function in neurons induces abnormal development of cerebral cortex and reactive gliosis in the brain. Genes Dev. 15, 859–876. 10.1101/gad.862101.

43. Gregorian, C., Nakashima, J., Le Belle, J., Ohab, J., Kim, R., Liu, A., Smith, K.B., Groszer, M., Garcia, A.D., Sofroniew, M.V., et al. (2009). Pten deletion in adult neural stem/progenitor cells enhances constitutive neurogenesis. J. Neurosci. Off. J. Soc. Neurosci. 29, 1874–1886. 10.1523/JNEUROSCI.3095-08.2009.

44. Tollefsen, S.E., and Kornfeld, R. (1983). The B4 lectin from Vicia villosa seeds interacts with N-acetylgalactosamine residues alpha-linked to serine or threonine residues in cell surface glycoproteins. J. Biol. Chem. 258, 5172–5176.

45. Baenziger, J.U., and Fiete, D. (1979). Structural determinants of concanavalin A specificity for oligosaccharides. J. Biol. Chem. 254, 2400–2407.

46. Kornfeld, R., and Ferris, C. (1975). Interaction of immunoglobulin glycopeptides with concanavalin A. J. Biol. Chem. 250, 2614–2619.

47. Tarentino, A.L., and Plummer, T.H. (1994). Enzymatic deglycosylation of asparagine-linked glycans: purification, properties, and specificity of oligosaccharide-cleaving enzymes from Flavobacterium meningosepticum. Methods Enzymol. 230, 44–57. 10.1016/0076-6879(94)30006-2.

48. Zawadzka, M., Rivers, L.E., Fancy, S.P.J., Zhao, C., Tripathi, R., Jamen, F., Young, K., Goncharevich, A., Pohl, H., Rizzi, M., et al. (2010). CNS-resident glial progenitor/stem cells produce Schwann cells as well as oligodendrocytes during repair of CNS demyelination. Cell Stem Cell 6, 578–590. 10.1016/j.stem.2010.04.002.

49. Howell, M.D., and Gottschall, P.E. (2012). Lectican proteoglycans, their cleaving metalloproteinases, and plasticity in the central nervous system extracellular microenvironment. Neuroscience 217, 6–18. 10.1016/j.neuroscience.2012.05.034.

50. Dolma, S., and Joshi, A. (2023). The Node of Ranvier as an Interface for Axo-Glial Interactions: Perturbation of Axo-Glial Interactions in Various Neurological Disorders. J. Neuroimmune Pharmacol. Off. J. Soc. NeuroImmune Pharmacol. 18, 215–234. 10.1007/s11481-023-10072-z.

51. Rasband, M.N., and Peles, E. (2021). Mechanisms of node of Ranvier assembly. Nat. Rev. Neurosci. 22, 7–20. 10.1038/s41583-020-00406-8.

52. Rios, J.C., Melendez-Vasquez, C.V., Einheber, S., Lustig, M., Grumet, M., Hemperly, J., Peles, E., and Salzer, J.L. (2000). Contactin-associated protein (Caspr) and contactin form a complex that is targeted to the paranodal junctions during myelination. J. Neurosci. Off. J. Soc. Neurosci. 20, 8354–8364. 10.1523/JNEUROSCI.20-22-08354.2000.

53. Li, C., Tropak, M.B., Gerlai, R., Clapoff, S., Abramow-Newerly, W., Trapp, B., Peterson, A., and Roder, J. (1994). Myelination in the absence of myelin-associated glycoprotein. Nature 369, 747–750. 10.1038/369747a0.

54. Marcus, J., Dupree, J.L., and Popko, B. (2002). Myelin-associated glycoprotein and myelin galactolipids stabilize developing axo-glial interactions. J. Cell Biol. 156, 567–577. 10.1083/jcb.200111047.

55. Yin, X., Crawford, T.O., Griffin, J.W., Tu, P. h, Lee, V.M., Li, C., Roder, J., and Trapp, B.D. (1998). Myelin-associated glycoprotein is a myelin signal that modulates the caliber of myelinated axons. J. Neurosci. Off. J. Soc. Neurosci. 18, 1953–1962. 10.1523/JNEUROSCI.18-06-01953.1998.

56. Schnaar, R.L., and Lopez, P.H.H. (2009). Myelin-associated glycoprotein and its axonal receptors. J. Neurosci. Res. 87, 3267–3276. 10.1002/jnr.21992.

57. Bull, C., Nason, R., Sun, L., Van Coillie, J., Madriz Sørensen, D., Moons, S.J., Yang, Z., Arbitman, S., Fernandes, S.M., Furukawa, S., et al. (2021). Probing the binding specificities of human Siglecs by cell-based glycan arrays. Proc. Natl. Acad. Sci. U. S. A. 118, e2026102118. 10.1073/pnas.2026102118.

58. Susuki, K., Chang, K.-J., Zollinger, D.R., Liu, Y., Ogawa, Y., Eshed-Eisenbach, Y., Dours-Zimmermann, M.T., Oses-Prieto, J.A., Burlingame, A.L., Seidenbecher, C.I., et al. (2013). Three mechanisms assemble central nervous system nodes of Ranvier. Neuron 78, 469–482. 10.1016/j.neuron.2013.03.005.

59. Arancibia-Carcamo, I.L., Ford, M.C., Cossell, L., Ishida, K., Tohyama, K., and Attwell, D. (2017). Node of Ranvier length as a potential regulator of myelinated axon conduction speed. eLife 6, e23329. 10.7554/eLife.23329.

60. Bradberry, M.M., Peters-Clarke, T.M., Shishkova, E., Chapman, E.R., and Coon, J.J. (2022). Highly fucosylated N-glycans at the synaptic vesicle and neuronal plasma membrane. Preprint at bioRxiv, 10.1101/2022.07.06.499060 10.1101/2022.07.06.499060.

61. Elbaz, B., Darwish, A., Vardy, M., Isaac, S., Tokars, H.M., Dzhashiashvili, Y., Korshunov, K., Prakriya, M., Eden, A., and Popko, B. (2024). The bone transcription factor Osterix controls extracellular matrix- and node of Ranvier-related gene expression in oligodendrocytes. Neuron 112, 247–263.e6. 10.1016/j.neuron.2023.10.008.

62. Recombinant lectin Gg for brain imaging of glycan structure and formation in the CNS node of Ranvier 10.1111/jnc.15695.

63. Luo, L., Ambrozkiewicz, M.C., Benseler, F., Chen, C., Dumontier, E., Falkner, S., Furlanis, E., Gomez, A.M., Hoshina, N., Huang, W.-H., et al. (2020). Optimizing Nervous System-Specific Gene Targeting with Cre Driver Lines: Prevalence of Germline Recombination and Influencing Factors. Neuron 106, 37–65.e5. 10.1016/j.neuron.2020.01.008.

64. Suttapitugsakul, S., Matsumoto, Y., Aryal, R.P., and Cummings, R.D. (2023). Large-Scale and Site-Specific Mapping of the Murine Brain *O* -Glycoproteome with IMPa. Anal. Chem. 95, 13423–13430. 10.1021/acs.analchem.3c00408.

65. Bekku, Y., Rauch, U., Ninomiya, Y., and Oohashi, T. (2009). Brevican distinctively assembles extracellular components at the large diameter nodes of Ranvier in the CNS. J. Neurochem. 108, 1266–1276. 10.1111/j.1471-4159.2009.05873.x.

66. Viapiano, M.S., Matthews, R.T., and Hockfield, S. (2003). A novel membrane-associated glycovariant of BEHAB/brevican is up-regulated during rat brain development and in a rat model of invasive glioma. J. Biol. Chem. 278, 33239–33247. 10.1074/jbc.M303480200.

67. Larsen, I.S.B., Narimatsu, Y., Joshi, H.J., Siukstaite, L., Harrison, O.J., Brasch, J., Goodman, K.M., Hansen, L., Shapiro, L., Honig, B., et al. (2017). Discovery of an O-mannosylation pathway selectively serving cadherins and protocadherins. Proc. Natl. Acad. Sci. U. S. A. 114, 11163–11168. 10.1073/pnas.1708319114.

68. van Vliet, S.J., van Liempt, E., Saeland, E., Aarnoudse, C.A., Appelmelk, B., Irimura, T., Geijtenbeek, T.B.H., Blixt, O., Alvarez, R., van Die, I., et al. (2005). Carbohydrate profiling reveals a distinctive role for the C-type lectin MGL in the recognition of helminth parasites and tumor antigens by dendritic cells. Int. Immunol. 17, 661–669. 10.1093/intimm/dxh246.

69. Ilarregui, J.M., Kooij, G., Rodrfguez, E., van der Pol, S.M.A., Koning, N., Kalay, H., van der Horst, J.C., van Vliet, S.J., Garcfa-Vallejo, J.J., de Vries, H.E., et al. (2019). Macrophage galactose-type lectin (MGL) is induced on M2 microglia and participates in the resolution phase of autoimmune neuroinflammation. J. Neuroinflammation 16, 130. 10.1186/s12974-019-1522-4.

70. Minami, A., Meguro, Y., Ishibashi, S., Ishii, A., Shiratori, M., Sai, S., Horii, Y., Shimizu, H., Fukumoto, H., Shimba, S., et al. (2017). Rapid regulation of sialidase activity in response to neural activity and sialic acid removal during memory processing in rat hippocampus. J. Biol. Chem. 292, 5645–5654. 10.1074/jbc.M116.764357.

71. Zeng, W.-F., Cao, W.-Q., Liu, M.-Q., He, S.-M., and Yang, P.-Y. (2021). Precise, fast and comprehensive analysis of intact glycopeptides and modified glycans with pGlyco3. Nat. Methods 18, 1515–1523. 10.1038/s41592-021-01306-0.

72. Watanabe, K., Taskesen, E., van Bochoven, A., and Posthuma, D. (2017). Functional mapping and annotation of genetic associations with FUMA. Nat. Commun. 8, 1826. 10.1038/s41467-017-01261-5.

