## Supplemental Material for "O-GalNAc glycans enrich in white matter tracts and regulate nodes of Ranvier"

Supplemental Figures 1-11

Supplemental Tables 1-2

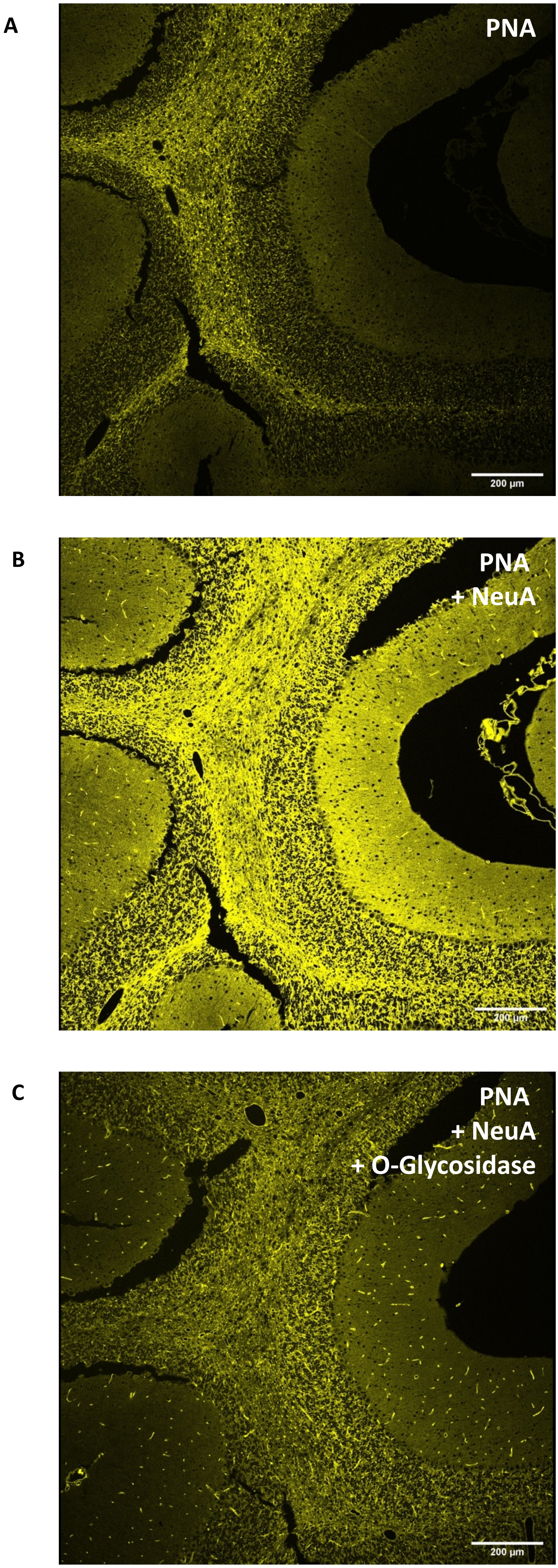


**Supplemental Figure 1. PNA binding in white matter tracts of the cerebellum. A)** PNA binding in coronal section of cerebellum were most intense signal in the arbor vitae. **B)** PNA binding after neuraminidase (NeuA) treatment broadly increased signal across the cerebellum. **C)** However, a portion of PNA binding following NeuA treatment was not sensitive to O-glycosidase, in particular within structures resembling microvasculature. Scale bar = 200 µm.

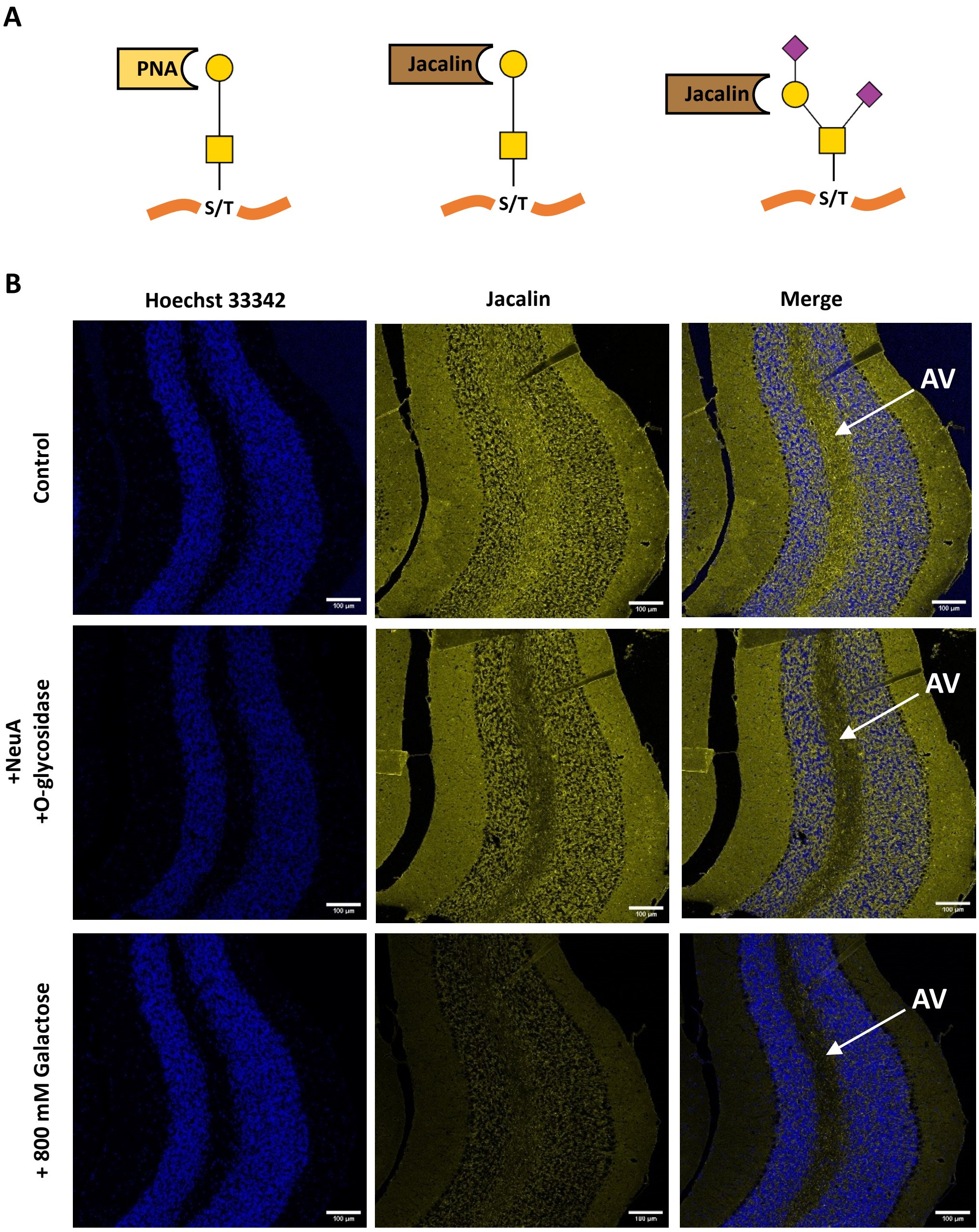


**Supplemental Figure 2. Jacalin binding to the arbor vitae was O-glycosidase sensitive, while other binding is not. A)** Binding preferences: PNA binds the galactose of O-GalNAc glycans only without sialic acid; Jacalin is reported to bind the galactose of O-GalNAc glycans independent of sialic acid. **B)** PNA and Jacalin binding to brain glycoproteins before and after O-glycosidase treatment and inhibition with 800 mM galactose. Jacalin binding is diffuse across all layers of the cerebellum, but only the signal in the arbor vitae is sensitive to O-glycosidase, consistent with O-glycans present in white matter tracks. High concentration of galactose reduced Jacalin binding across all layers, though some residual signal was detected. White matter tracts of the arbor vitae (AV) are indicated with white arrows in the right column for reference. Scale bars = 100 µm.

A

B

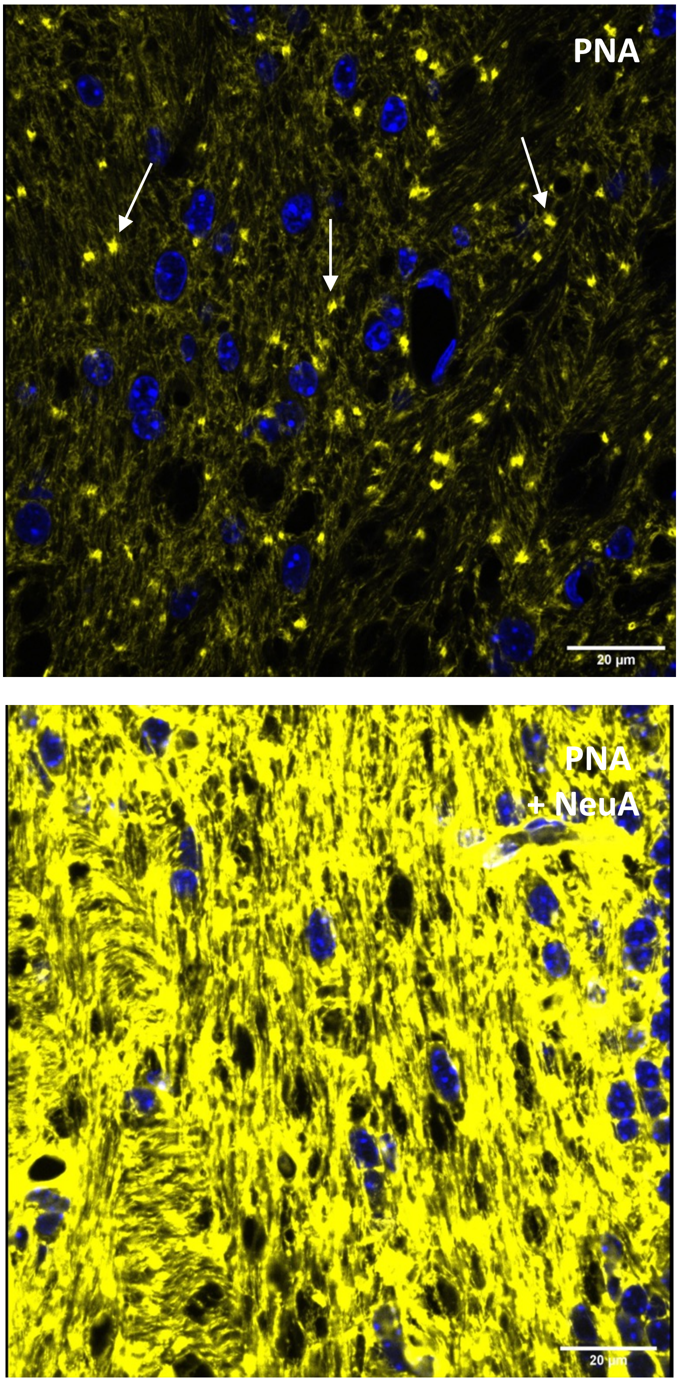


**Supplemental Figure 3. Punctate PNA binding in cerebellar white matter. A)** High magnification of PNA binding in coronal section of cerebellum showed several punctate structures with high intensity (white arrows). **B)** PNA binding after neuraminidase (NeuA) showed a more diffuse pattern and brighter signal. Scale bar = 20 µm.

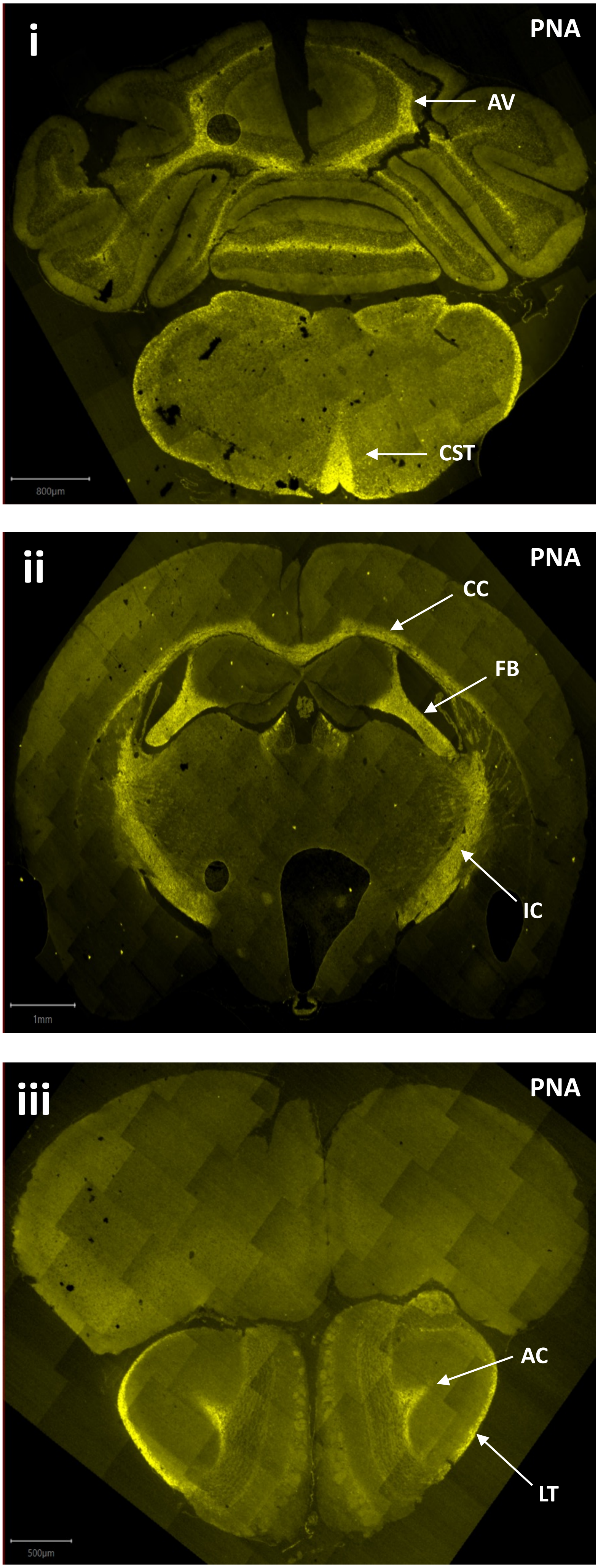


**Supplemental Figure 4. PNA binding enriches in multiple key white matter tracts across the brain.** PNA preferentially binds galactose of O-GalNAc glycans lacking sialic acid. Across the entirety of the brain PNA binding is highest in white matter structures. Posteriorly **(i)**, high binding is noted in the arbor vitae (AV) and corticospinal tract (CST) of the brain stem pyramids. Coronal sections including diencephalic structures **(ii)** are notable for strong binding in white matter structures including the corpus collosum (CC), fimbria (FB), and internal capsule (IC). Anteriorly **(iii)**, strong PNA binding is seen in white matter tracts including the anterior commissure (AC) and lateral tract (LT) of the olfactory bulb. Scale bars i = 800 µm, ii = 1 mm, iii = 500 µm.

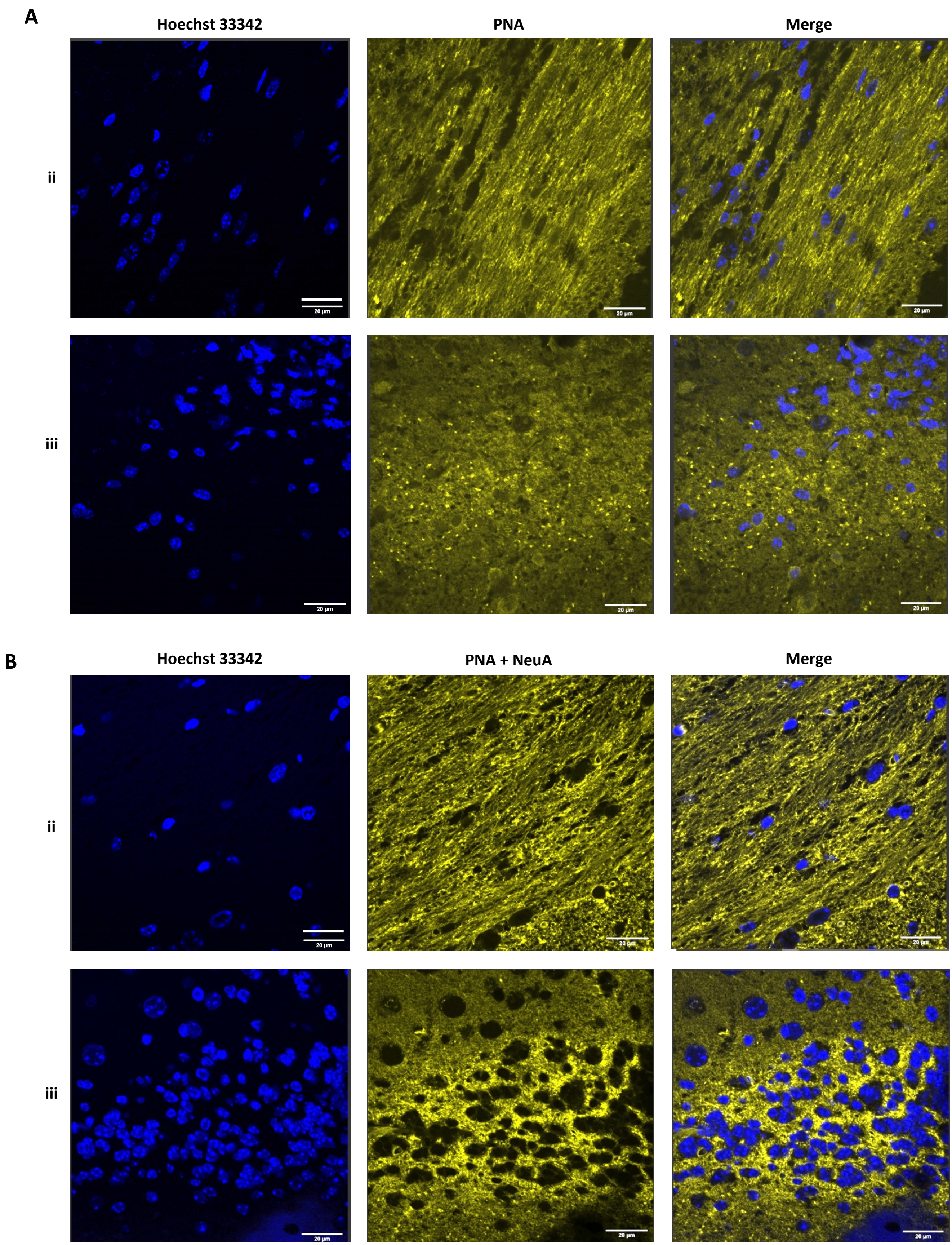


**Supplemental Figure 5. PNA binding in white matter tracts differs based on orientation of fibers. A)** PNA binding in coronal section of the corpus collosum (iI), where the majority of white matter tracts run parallel with the section display a striated pattern. In contrast, white matter tracts of the olfactory bulb (iii) which run orthogonal to the coronal section display a more discrete and punctate pattern. **B)** PNA binding after neuraminidase treatment in the corpus collosum (ii) and olfactory bulb (iii) show a similar pattern. Coronal sections (ii, iii) correspond to locations indicated in Fig. 2D. Scale bar = 20 µm.

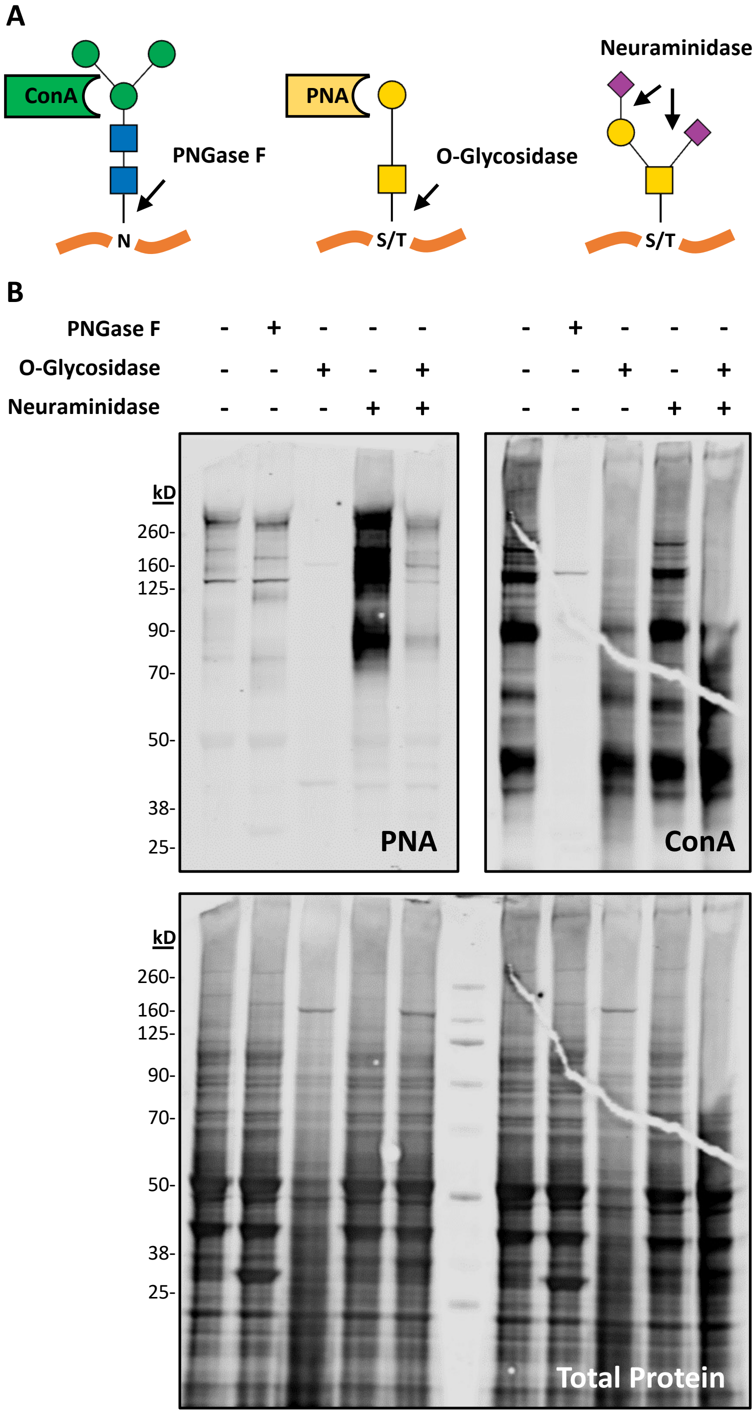


**Supplemental Figure 6. Lectin blotting of brain lysate confirms the presence of highly sialylated O-glycans.** **A)** Binding preferences and glycosidase sensitivity: ConA binds the core structure of N-glycans and is cleaved by PNGase F; PNA binds T-antigen and is cleaved by O-glycosidase; Neuraminidase cleaves sialic acid in all linkages **B)** PNA and ConA binding to brain glycoproteins before and after glycosidase treatments. PNA signal is present but low at baseline conditions, is insensitive to PNGase F, but reduced by O-glycosidase. Neuraminidase treatment results in a dramatic increase in PNA binding which is reduced by O-glycosidase, consistent with the majority of brain O-glycans containing sialic acid. ConA signal is only sensitive to PNGase F treatment. Each lane contains 15 µg of cortical lysates precleared with streptavidin beads, treated with or without the indicated glycosidases for 1 hour at 37°C according to manufacturer's protocol. Total protein stain is shown below for the full blot, which was subsequently cut in half for the different lectin blotting.

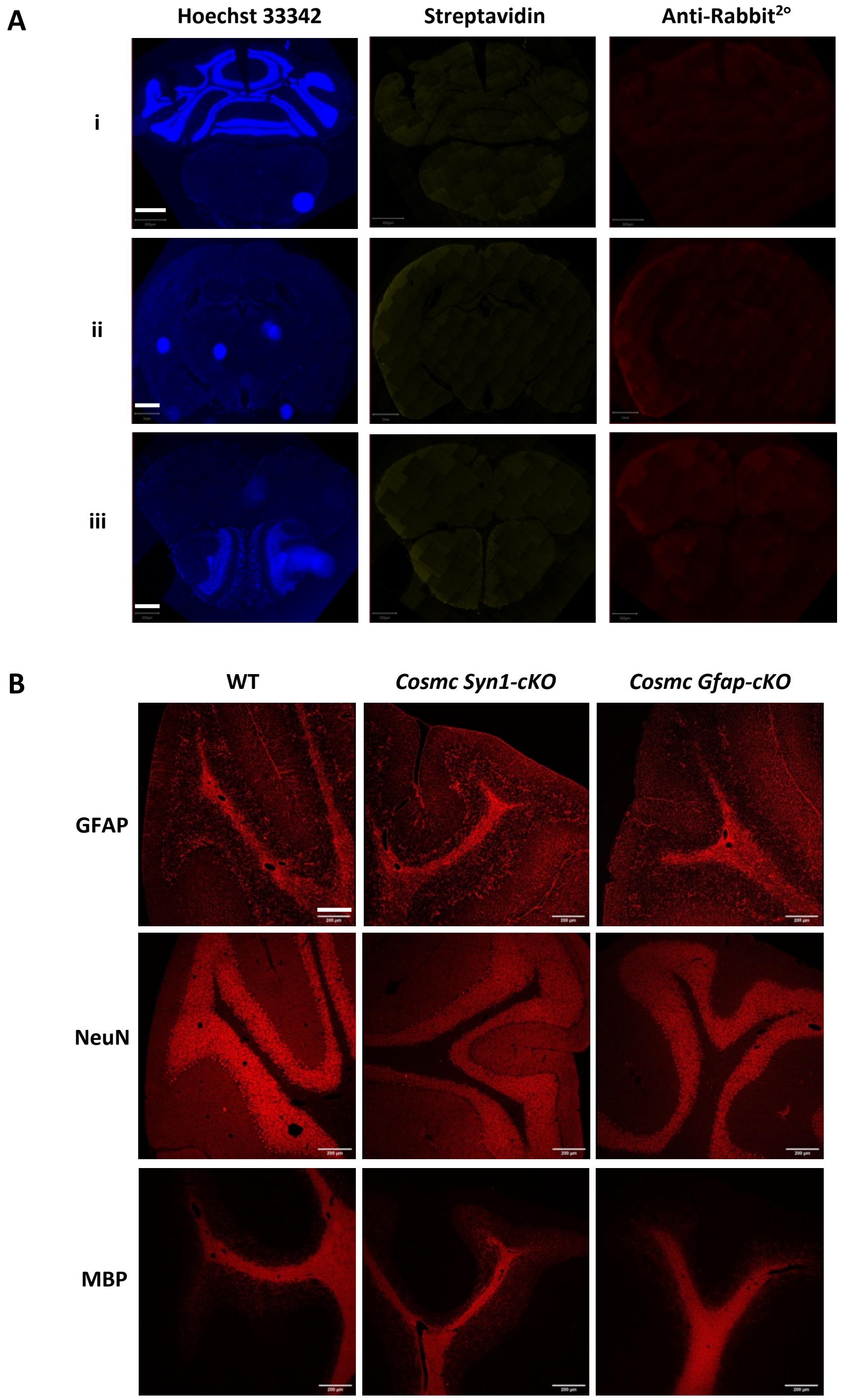


**Supplemental Figure 7. Lectin/antibody controls and gross morphology of *Cosmc Syn1- and Gfap cKO* lines. A)** Control binding of FITC conjugated streptavidin and secondary antibody alone (anti-rabbit 588) showed minimal background in wildtype mice. Coronal sections (i, ii, iii) correspond to locations indicated in Fig. 2D. Scale bars i = 800 µm, ii = 1 mm, iii = 500 µm. **B)** Cerebellar morphology assessed by staining with common markers including GFAP, NeuN, and MBP, is unchanged in *Cosmc* cKO lines. Scale bar = 200 µm.

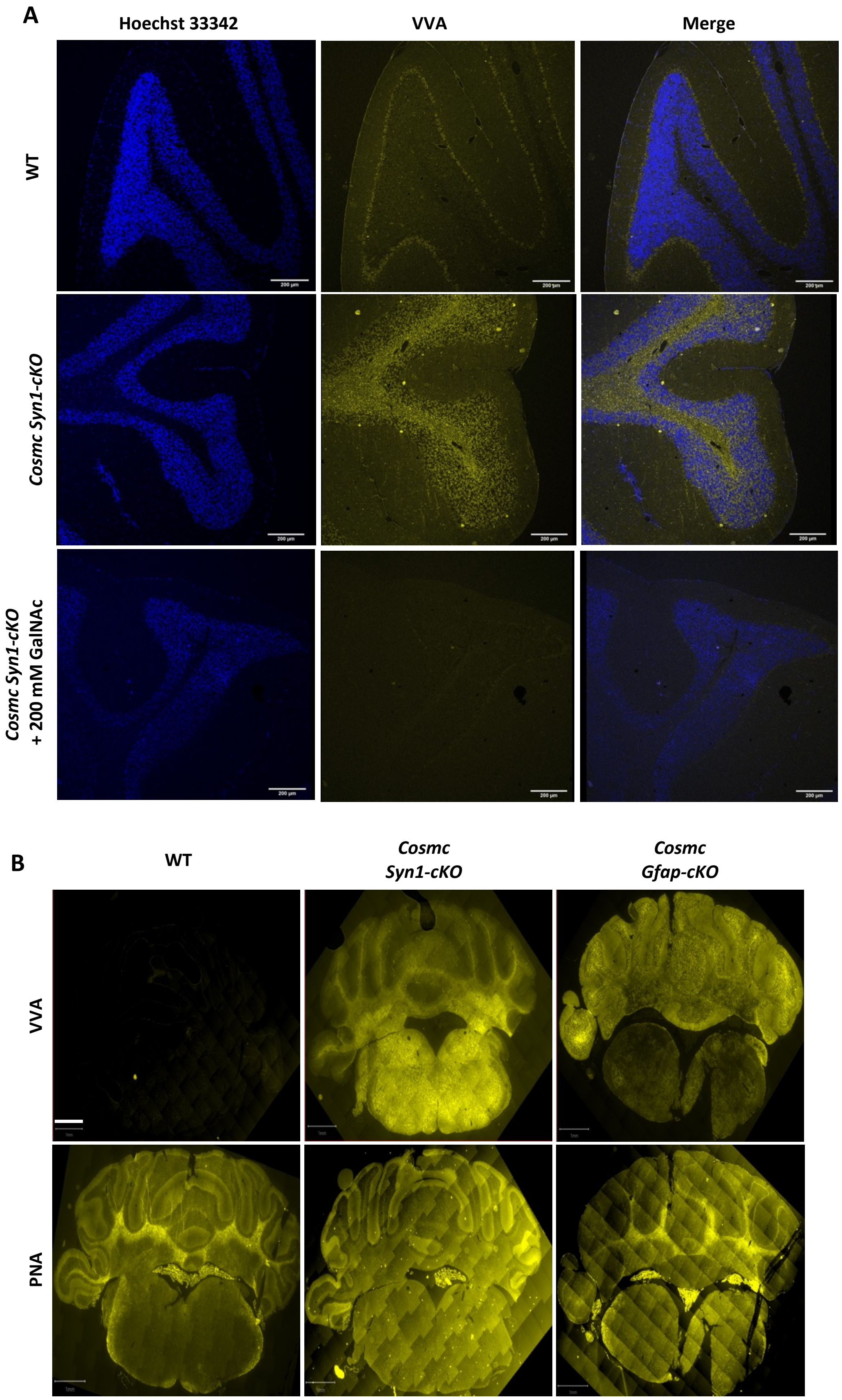


**Supplemental Figure 8. VVA specificity and VVA/PNA binding in cerebellum of *Cosmc cKO* lines.** **A)** VVA binding across the cerebellum is absent in WT mice. VVA binding in the arbor vitae of *Cosmc Syn1 cKO* mice is blocked by incubation with 200 mM GalNAc. Scale bar = 200 µm. **B)** VVA and PNA binding in the cerebellum of *Cosmc cKO* lines. Scale bar = 1 mm.

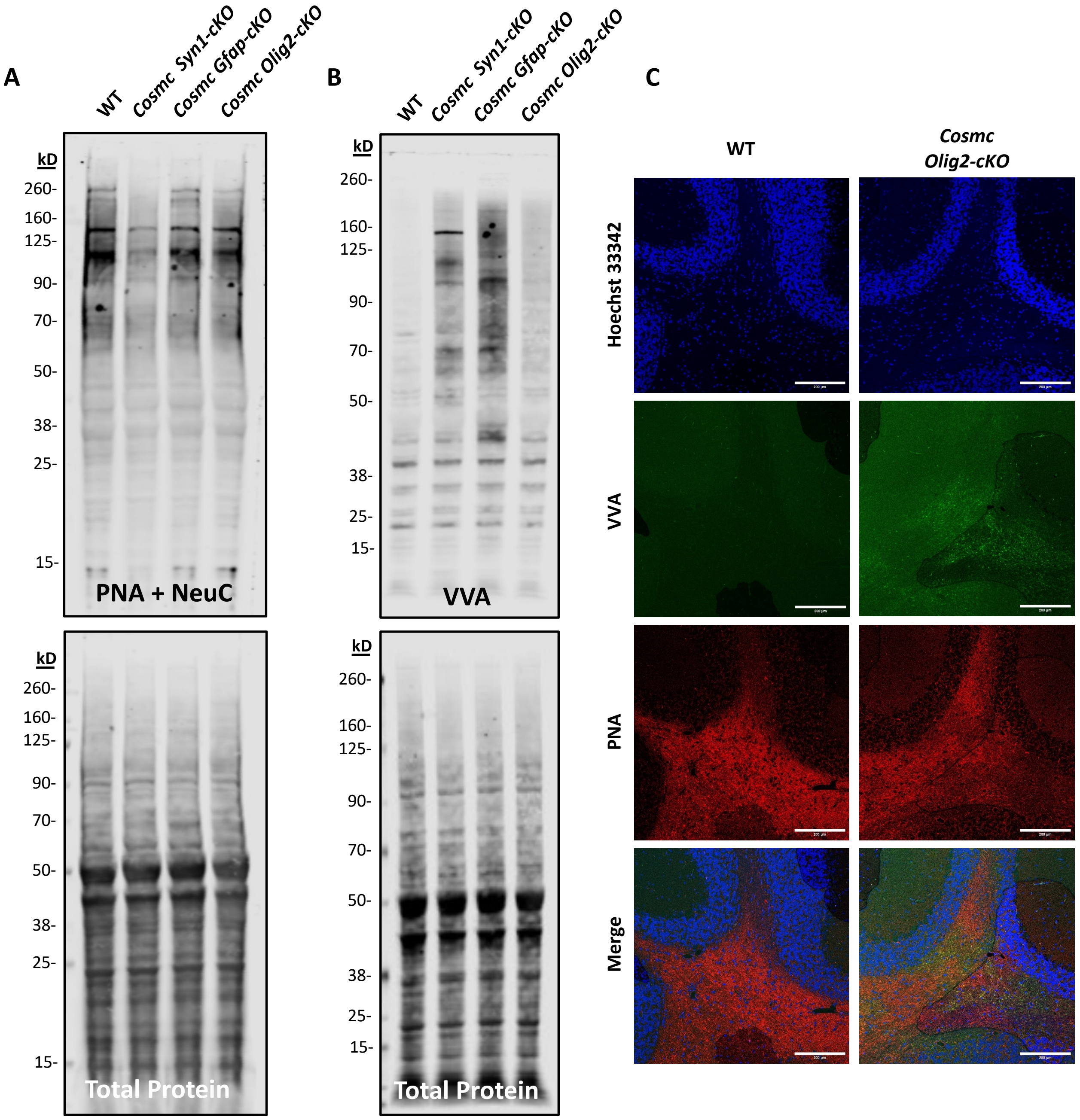


**Supplemental Figure 9. Genetic deletion of *Cosmc* in *Olig2* expressing cells had minimal effect on PNA and VVA signal in brain**. **A)** Total levels of O-GalNAc glycans detected by PNA blotting after neuraminidase treatment showed no clear change in the cortex of the C*osmc Olig2-cKO* line, similar to C*osmc-Syn1* and C*osmc Gfap-cKO* lines. Total protein stain shown in lower panel. **B)** Low levels of VVA binding to Tn-antigen were detected in the cortex of the *Cosmc Olig2-cKO* line, similar to wild-type brain and far less than C*osmc Syn1* and C*osmc Gfap-cKO* lines, suggesting a smaller contribution of these cell lineages to O-GalNAc synthesis in the brain. Total protein stain shown in lower panel. **C)** VVA and PNA binding in the cerebellum of *Cosmc Olig2-cKO* lines. Scale bar = 200 µm.

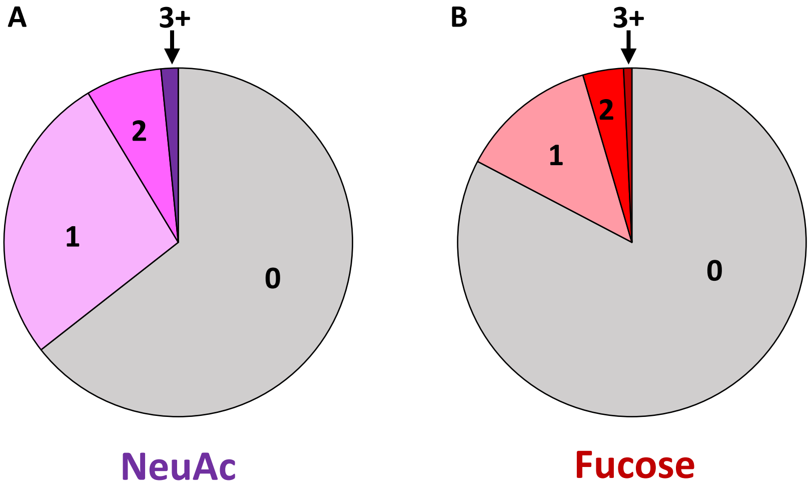


**Supplemental Figure 10. LC-MS analysis of brain O-GalNAc glycans after neuraminidase is consistent with a partial desialylation reaction. A)** In comparison to native cortex analyzed by MALDI-MS, where over 90 % of O-GalNAc glycans are sialylated (Williams, *et al.*, 2022), LC-MS/MS analysis of O-GalNAc glycans enriched by PNA binding after neuraminidase treatment revealed that ~35% of structures were sialylated. The majority of structures contained 1 sialic acid, suggesting that a partial desialylation reaction occurred, as in the untreated brain the disialylated T-antigen makes up nearly 70% of the total signal. **B)** LC-MS/MS analysis of O-GalNAc glycans using the same protocol revealed that majority of structures (~83%) lacked fucose, consistent with prior MALDI-MS data.

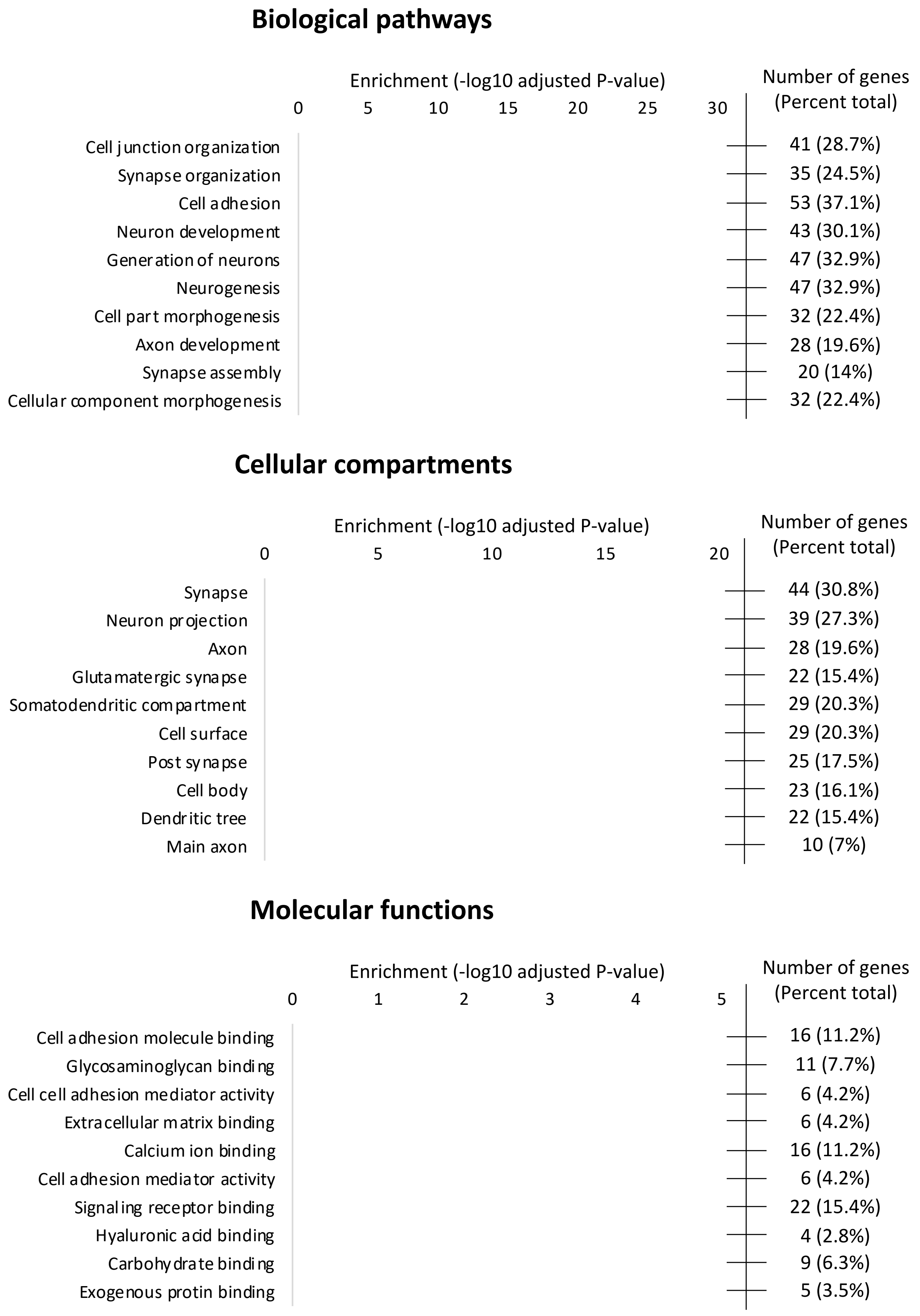


**Supplemental Figure 11. Gene Ontology (GO) analysis of O-GalNAc containing proteins. A)** GO analysis of biological pathways shows significant enrichment for genes involved with the synapse, cell adhesion, and development. **B)** GO analysis of cellular compartments shows significant enrichment for genes expressed in synapse, neurons, and axon. **C)** GO analysis of molecular functions shows enrichment for genes involved in cell adhesion, glycosaminoglycan binding, and the extracellular matrix. Data was analyzed using the FUMA GENE2FUNC platform, with the top 10 findings from each category presented.

**A**

**FRONTAL CORTEX**

| Cluster | *Galnt1* | *Galnt3* | *Galnt4* | *Galnt6* | *Galnt7* | *Galnt9* | *Galnt10* | *Galnt11* | *Galnt12* | *Galnt13* | *Galnt14* | *Galnt15* | *Galnt16* | *Galnt18* | *Galntl6* | Sum |
|---|---|---|---|---|---|---|---|---|---|---|---|---|---|---|---|---|
| Interneuron_CGE_Cplx3-Synpr [#1] | 6 | 1 | 1 | 1 | 5 | 3 | 2 | 4 | 1 | 6 | 9 | 1 | 5 | 4 | 1 | **50** |
| Interneuron_MGE_Sst-Pvalb [#2] | 7 | 1 | 1 | 1 | 8 | 6 | 2 | 4 | 1 | 4 | 4 | 1 | 7 | 2 | 2 | **51** |
| Neuron_Layer6Subplate_Syt6 [#3] | 5 | 1 | 1 | 1 | 2 | 21 | 1 | 4 | 1 | 5 | 1 | 1 | 11 | 4 | 1 | **60** |
| Neuron_Layer5b_Fezf2 [#4] | 7 | 1 | 1 | 1 | 5 | 11 | 1 | 4 | 1 | 9 | 6 | 1 | 8 | 5 | 1 | **62** |
| Neuron_Claustrum_Nr4a2 [#5] | 3 | 1 | 1 | 1 | 3 | 2 | 1 | 4 | 1 | 4 | 15 | 1 | 5 | 4 | 1 | **47** |
| Neuron_Layer23_Nptxr [#6] | 4 | 1 | 1 | 1 | 5 | 7 | 1 | 4 | 1 | 5 | 3 | 1 | 10 | 4 | 1 | **49** |
| Neuron_Layer5_Parm1 [#7] | 5 | 1 | 1 | 1 | 5 | 25 | 1 | 3 | 1 | 11 | 3 | 1 | 22 | 4 | 1 | **85** |
| Astrocyte_Gja1 [#8] | 11 | 1 | 2 | 1 | 4 | 2 | 1 | 3 | 1 | 2 | 1 | 1 | 7 | 4 | 1 | **42** |
| Oligodendrocyte_Tfr [#9] | 7 | 1 | 1 | 27 | 5 | 2 | 1 | 3 | 1 | 2 | 1 | 1 | 4 | 1 | 1 | **58** |
| Polydendrocyte_Tnr [#10] | 11 | 3 | 1 | 2 | 3 | 2 | 9 | 3 | 1 | 8 | 1 | 1 | 6 | 1 | 1 | **53** |
| Microglia_Macrophage_C1qb [#11] | 14 | 2 | 2 | 1 | 5 | 2 | 4 | 3 | 4 | 4 | 1 | 1 | 4 | 2 | 1 | **50** |
| Endothelial_Flt1 [#12] | 9 | 1 | 2 | 1 | 4 | 2 | 2 | 3 | 1 | 2 | 1 | 4 | 2 | 4 | 1 | **39** |
| Mural_Rgs5Acta2 [#13] | 15 | 1 | 2 | 1 | 4 | 2 | 2 | 3 | 1 | 2 | 1 | 2 | 3 | 4 | 1 | **44** |
| Fibroblast-Like_Dcn [#14] | 11 | 1 | 2 | 1 | 3 | 2 | 2 | 4 | 2 | 6 | 1 | 1 | 2 | 2 | 1 | **41** |

**B**

**CEREBELLUM**

| Cluster | *Galnt1* | *Galnt4* | *Galnt6* | *Galnt7* | *Galnt9* | *Galnt10* | *Galnt11* | *Galnt12* | *Galnt13* | *Galnt14* | *Galnt15* | *Galnt16* | *Galnt18* | *Sum* |
|---|---|---|---|---|---|---|---|---|---|---|---|---|---|---|
| GranularNeuron_Gabra6 [#1] | 4 | 1 | 1 | 8 | 11 | 1 | 4 | 1 | 33 | 1 | 4 | 2 | 2 | 73 |
| PurkinjeNeuron_Pcp2 [#2] | 6 | 1 | 1 | 3 | 9 | 1 | 3 | 1 | 23 | 1 | 1 | 3 | 1 | 54 |
| Interneurons_Pvalb [#3] | 6 | 1 | 1 | 9 | 6 | 1 | 7 | 1 | 18 | 2 | 1 | 5 | 14 | 72 |
| Interneurons_and_Other_Nnat [#4] | 5 | 1 | 1 | 7 | 2 | 1 | 5 | 1 | 8 | 2 | 2 | 4 | 2 | 41 |
| Microglia_Macrophage_C1qb [#5] | 11 | 3 | 2 | 4 | 4 | 1 | 1 | 4 | 1 | 1 | 1 | 1 | 1 | 35 |
| Oligodendrocyte_Polydendrocyte_Tfr_Tnr [#6] | 6 | 1 | 23 | 7 | 1 | 3 | 4 | 1 | 4 | 1 | 1 | 4 | 2 | 58 |
| BergmannGlia_Gpr37l1 [#7] | 9 | 2 | 1 | 6 | 1 | 2 | 3 | 1 | 3 | 1 | 1 | 7 | 2 | 39 |
| Astrocyte_Gja1 [#8] | 9 | 2 | 1 | 5 | 1 | 6 | 6 | 1 | 4 | 1 | 1 | 4 | 10 | 51 |
| Choroid_Plexus_Ttr [#9] | 5 | 4 | 1 | 2 | 1 | 3 | 5 | 1 | 1 | 1 | 2 | 1 | 3 | 30 |
| Endothelial_Flt1 [#10] | 6 | 3 | 1 | 5 | 2 | 2 | 3 | 1 | 3 | 1 | 6 | 1 | 5 | 39 |
| Fibroblast-Like_Dcn [#11] | 11 | 1 | 1 | 4 | 2 | 4 | 3 | 2 | 2 | 1 | 1 | 1 | 5 | 38 |

**C**

**FRONTAL CORTEX**

| Cluster | *C1galt1* | *C1galt1c1* | *Gfap* | *Syn1* | *Olig2* |
|---|---|---|---|---|---|
| Interneuron_CGE_Cplx3-Synpr [#1] | 2 | 5 | 1 | 25 | 1 |
| Interneuron_MGE_Sst-Pvalb [#2] | 1 | 4 | 1 | 29 | 1 |
| Neuron_Layer6Subplate_Syt6 [#3] | 1 | 3 | 1 | 41 | 1 |
| Neuron_Layer5b_Fezf2 [#4] | 2 | 3 | 1 | 25 | 1 |
| Neuron_Claustrum_Nr4a2 [#5] | 2 | 4 | 1 | 47 | 1 |
| Neuron_Layer23_Nptxr [#6] | 2 | 3 | 1 | 40 | 1 |
| Neuron_Layer5_Parm1 [#7] | 2 | 2 | 1 | 27 | 1 |
| Astrocyte_Gja1 [#8] | 1 | 5 | 23 | 5 | 6 |
| Oligodendrocyte_Tfr [#9] | 2 | 7 | 1 | 6 | 23 |
| Polydendrocyte_Tnr [#10] | 2 | 6 | 1 | 8 | 78 |
| Microglia_Macrophage_C1qb [#11] | 2 | 5 | 1 | 11 | 2 |
| Endothelial_Flt1 [#12] | 2 | 5 | 1 | 5 | 1 |
| Mural_Rgs5Acta2 [#13] | 1 | 5 | 1 | 6 | 1 |
| Fibroblast-Like_Dcn [#14] | 1 | 6 | 2 | 8 | 2 |

**D**

**CEREBELLUM**

| Cluster | *C1galt1* | *C1galt1c1* | *Gfap* | *Syn1* | *Olig2* |
|---|---|---|---|---|---|
| GranularNeuron_Gabra6 [#1] | 1 | 7 | 1 | 16 | 1 |
| PurkinjeNeuron_Pcp2 [#2] | 1 | 3 | 1 | 9 | 1 |
| Interneurons_Pvalb [#3] | 1 | 5 | 1 | 20 | 1 |
| Interneurons_and_Other_Nnat [#4] | 1 | 5 | 1 | 20 | 1 |
| Microglia_Macrophage_C1qb [#5] | 1 | 3 | 1 | 1 | 1 |
| Oligodendrocyte_Polydendrocyte_Tfr_Tnr [#6] | 2 | 5 | 2 | 2 | 30 |
| BergmannGlia_Gpr37l1 [#7] | 1 | 11 | 5 | 2 | 1 |
| Astrocyte_Gja1 [#8] | 1 | 4 | 46 | 3 | 1 |
| Choroid_Plexus_Ttr [#9] | 2 | 8 | 6 | 1 | 1 |
| Endothelial_Flt1 [#10] | 1 | 6 | 1 | 2 | 1 |
| Fibroblast-Like_Dcn [#11] | 2 | 6 | 1 | 2 | 1 |

**Supplemental Table 1. Single cell expression data from mouse brain of genes involved in the first two steps of O-GalNAc Synthesis.** Data downloaded from www.dropviz.org. Values represent transcripts per 100,000 in a cluster, generated obtaining the raw data as the natural log transformation and corrected using the exponential function. Of note, transcripts that equal zero in the raw data become 1 after exponential transformation, limiting the quantitative interpretation of low abundance transcripts. *Galnt* levels across the frontal cortex **(A)** and cerebellum **(B)** are shown, including a sum of all detected *Galnts*. *Galnts* not detected across any of the clusters were omitted. Levels of T-Synthase (*C1galt1*), Cosmc (*C1galt1c1*), *Gfap*, and *Syn1* from frontal cortex **(C)** and cerebellum **(D)**.

| Genotype | ***Cosmc-Syn1-cKO*** | | | ***Cosmc-Gfap-cKO*** | | | ***Cosmc-Olig2-cKO*** | | |
|---|---|---|---|---|---|---|---|---|---|
| **Test** | Fisher's exact test | | | Fisher's exact test | | | Fisher's exact test | | |
| **P value** | 0.721 | | | >0.9999 | | | >0.9999 | | |
| **P value summary** | ns | | | ns | | | ns | | |
| **One- or two-sided** | Two-sided | | | Two-sided | | | Two-sided | | |
| **Statistically significant (P < 0.05)?** | No | | | No | | | No | | |
| **Data analyzed** | Male | Female | Total | Male | Female | Total | Male | Female | Total |
| **WT** | 13 | 11 | 24 | 8 | 6 | 14 | 3 | 4 | 7 |
| **KO** | 7 | 4 | 11 | 9 | 7 | 16 | 4 | 6 | 10 |
| **Total** | 20 | 15 | 35 | 17 | 13 | 30 | 7 | 10 | 17 |
| **Percentage of row total** | Male | Female | | Male | Female | | Male | Female | |
| **WT** | 54.17% | 45.83% | | 57.14% | 42.86% | | 42.86% | 57.14% | |
| **KO** | 63.64% | 36.36% | | 56.25% | 43.75% | | 40.00% | 60.00% | |
| **Percentage of column total** | Male | Female | | Male | Female | | Male | Female | |
| **WT** | 65.00% | 73.33% | | 47.06% | 46.15% | | 42.86% | 40.00% | |
| **KO** | 35.00% | 26.67% | | 52.94% | 53.85% | | 57.14% | 60.00% | |
| **Percentage of grand total** | Male | Female | | Male | Female | | Male | Female | |
| **WT** | 37.14% | 31.43% | | 26.67% | 20.00% | | 17.65% | 23.53% | |
| **KO** | 20.00% | 11.43% | | 30.00% | 23.33% | | 23.53% | 35.29% | |

**Supplemental Table 2. Birth ratios between male and females show no significant difference between genotypes**
